# Adversarial interspecies relationships facilitate population suppression by gene drive in spatially explicit models

**DOI:** 10.1101/2022.05.08.491087

**Authors:** Yiran Liu, WeiJian Teo, Haochen Yang, Jackson Champer

## Abstract

Suppression gene drives are designed to bias their inheritance and increase in frequency in a population, disrupting an essential gene in the process. When the frequency is high enough, the population will be unable to reproduce above the replacement level and could be eliminated. CRISPR suppression drives based on the homing mechanism have already seen success in the laboratory, particularly in malaria mosquitoes. However, several models predict that the use of these drives in realistic populations with spatial structure may not achieve complete success. This is due to the ability of wild-type individuals to escape the drive and reach empty areas with reduced competition, allowing them to achieve high reproductive success and leading to extinction-recolonization cycles across the landscape. Here, we extend our continuous space gene drive framework to include two competing species or predator-prey species pairs. We find that in both general and mosquito-specific models, the presence of a competing species or predator can greatly facilitate drive-based suppression, even for drives with modest efficiency. However, the presence of a competing species also substantially increases the frequency of outcomes in which the drive is lost before suppression is achieved. These results are robust in models with seasonal population fluctuations that mosquito populations often experience. We also found that suppression can be somewhat more difficult if targeting a predator with strong predator-prey interactions. Our results illustrate the difficulty of predicting outcomes of interventions that could substantially affect populations of interacting species in complex ecosystems. However, our results are also potentially promising for the prospects of less powerful suppression gene drives for achieving successful elimination of target mosquito and other pest populations.

## Introduction

Experiments in mosquitoes^1–5^ and flies^6,7^ have demonstrated that homing suppression gene drives could potentially be developed in many organisms. These engineered alleles are designed to spread through a population while targeting a haplosufficient but essential gene, eventually causing population suppression when they reach high frequency in a population^8–12^. The most successful example thus far has been a CRISPR drive construct targeting *doublesex* in *Anopheles gambiae* malaria mosquitoes, spreading rapidly and inducing suppression in just a few generations^2^. In such homing drives, the guide RNA (gRNA) directs Cas9 to cut a wild-type chromosome at the target gene in the germline. This wild-type allele then undergoes homology-directed repair (HDR), copying the drive allele to the target site and thus converting the wildtype allele to a drive allele. The drive allele is then inherited at an increased rate, allowing it to increase in frequency in the population.

However, if end-joining repair rather than HDR occurs, the wild-type site might mutate into a resistance allele, which cannot be converted to a desirable allele anymore^13^. These resistance alleles might slow down the drive if they result in a nonfunctional target gene, but if they retain target gene function, they will usually cause the suppression drive fail due to their large advantage over drive alleles. Targeting of highly conserved genomic sites can reduce the chance that functional resistance alleles form^2^, but this strategy is not always effective^5^. Multiplexing of gRNAs targeting adjacent sites can potentially be quite effective at any target site^6,14^, but this comes at the cost of drive conversion efficiency for more than perhaps 2-4 gRNAs^14^. If drive conversion efficiency is too low, the suppression drive may lack the power to suppress the population, resulting in an equilibrium drive allele frequency and a somewhat reduced population, depending on drive and population characteristics.

Compared to panmictic population models, potentially more realistic spatial models showed that even higher efficiency is necessary to achieve a successful suppression outcome. In models where individuals move over in a continuous space, the “chasing” phenomenon might occur after the release of a suppression drive^15^. In this situation, wild-type individuals might benefit from the lack of competition in empty areas cleared by the drive, allowing them to quickly reproduce before the drive arrive again from an adjacent region. The drive and wild-type alleles could coexist until the end of simulation, with the drive “chasing” the wild-type alleles but never eliminating them all^15–20^. To avoid this undesirable state and quickly eliminate a population, higher overall drive quality is required^15,17^. Drive conversion efficiency is sufficient in *Anopheles gambiae*, but fitness costs due to leaky somatic Cas9 expression would still possibly prevent population elimination in spatial models^20^. In *Drosophila melanogaster*, drive conversion efficiencies are generally lower, and embryo resistance allele formation is higher, even though somatic fitness costs are potentially less of an issue^6^. Drive conversion efficiency is even lower still in mice^21,22^ and *Aedes aegypt*^23–25^. Thus, prospects for successful elimination of target populations are not favorable in current models, at least with existing drive components.

However, increasing the complexity of gene drives models has often resulted in substantially different results compared to simplified models, as exemplified by the chasing phenomenon emerging in continuous space models. A further increase in complexity could involve increased ecological fidelity, modeling interspecies interactions. We hypothesize that such interactions could add additional pressure to a species targeted by a suppression drive, potentially easing the difficulty of achieving a successful result. For example, other species might share a partially overlapping ecological niche or directly prey on the target species. These factors could prevent a heavily suppressed population from achieving their highest potential “intrinsic growth rate” at low densities despite low population numbers in a chasing environment.

Models have previously used relatively high values for low-density growth rates based on observed rapid population increases at the beginning of wet seasons in regions that are highly seasonal^15,26^. However, it is likely that competitors and predators would start at similarly low density at the beginning of a wet season, preventing them from substantially influencing growth at this time. Later in the rainy season, they might be at higher densities that prevent the target species from achieving the same low-density growth rate that was possible earlier in the season. Other existing models are based on intrinsic species characteristics and observed introductions rather than observations of the species in environments with potential competitors or predators^19^. Thus, multi-species models may be necessary for accurately modeling the population dynamics of a species when the normal ecological balance is altered by a gene drive or other phenomena that causes a sudden population change.

Here, we simulate suppression gene drives in spatial models with more complex ecological environments that include both a target species and another interacting species. Specifically, we investigate how competition and predation relationships might affect the outcomes of a suppression strategy in both simple discrete-generation and mosquito-specific models. Overall, we find that the increased pressure from competitors or predators can substantially increase the chance of success for drives with even moderate efficiency.

## Methods

### Gene drive strategy

The suppression drives in this study are based on targeting a haplosufficient but essential female fertility gene. In this gene drive, CRISPR/Cas9 is used to cut the wild-type gene as directed by one or more guide RNAs (gRNAs). The target chromosome then undergoes homology-directed repair, resulted the drive allele being copied to the target site (“drive conversion”). Female drive heterozygotes still are fertile due to the target gene haplosufficiency. However, when the drive allele eventually reaches high frequency, increased rates of sterile drive homozygous females will result in decreased total population size or even population collapse.

Instead of homology-directed repair, if the cleaved target site undergoes end-joining repair, this will cause the target site to mutate into a resistance allele, which cannot be recognized by Cas9 for future cleavage, preventing successful drive conversion. If this happens, the target gene will most often be rendered nonfunctional, and the allele will be called a nonfunctional resistance allele (often called “r2 alleles”). Such alleles can still cause female sterility when homozygous or when paired with drive alleles. Thus, they often will not prevent population suppression, but they can still slow down the spread of the gene drive and reduce its suppressive power. A more drastic outcome occurs if the target gene is mutated into a functional resistance allele (“r1 allele”), which still allows female fertility. The presence of such an allele will usually cause immediate failure of suppression drive, with the population quickly rebounding. Experimental studies have shown that targeting a highly conserved site or using multiple gRNAs could help reducing the occurrence of “r1” resistance. Though the former strategy is often insufficient alone, and the latter strategy leads to reduced efficiency with high numbers of gRNAs, these methods can be combined for even higher reduction of functional resistance. Furthermore, reducing such alleles to an acceptably low level is likely a prerequisite for releasing a suppression drive into a population. Accordingly, we only consider nonfunctional resistance in this study (though see our previous studies for an assessment of functional resistance alleles for homing drives in spatially continuous models^15,17,19^).

### Simulation model

This model is based on the previous studies with one exception noted below^15,17,20^. In short, we simulated sexually reproducing diploids with the forward genetic simulation software SLiM (version 3.7)^27^ with 50,000 population size. The initial wild-type population is allowed to reproduce for 10 generations to achieve an equilibrium before gene drive individuals are dropped into the center of population.

The whole gene drive process occurs independently in each stage of each reproducing individual. Wild-type alleles in drive heterozygous parents are converted to a drive allele with a probability equal to drive efficiency (also called drive conversion rate). The wild-type allele also has a probability to mutate into a resistance allele equal to the “germline resistance allele formation rate”, which we set half of (1 – drive conversion rate) as default. If offspring contain any wild-type alleles and the mother has a drive allele, then the wild-type alleles may be mutated into resistance alleles due to maternally deposited Cas9/gRNA activity in the early embryo. Our default parameters represent a relatively efficient drive (somewhat better than existing fly designs^6^, though still substantially less efficient drive conversion than existing *Anopheles* drives^2^) with 80% drive conversion rate and 10% embryo resistance allele formation rate.

There are also two kinds of fitness costs that might affect our suppression drive. One is direct fitness, which we define for drive homozygous individuals relative to wild-type individuals, with multiplicative fitness for each drive allele. The default value for this direct fitness cost is 0.95, indicating only small reductions in fitness compared to wild-type individuals (with a fitness of 1) due to the cost of expression of all drive allele elements. Another fitness cost comes specifically from undesired somatic expression in female heterozygotes, which could disrupt wild-type alleles in somatic cells, causing reduced female fertility in only drive/wild-type heterozygotes. The default for this parameter is 0.9, indicating some leaky somatic Cas9 expression, less than observed in *Anopheles* studies but perhaps more than seen with the *nanos* promoter in *Drosophila*. These fitness reductions can have different effects, depending on the model.

### Discrete-generation model

We created a model with discrete, non-overlapping generations. We tracked the position of each individual across a 1×1 unitless arena. 50,000 wild-type individuals were randomly placed into the arena, and 500 gene drive heterozygotes were released after 10 generations. In this model, the competition strength between individuals depends on distance where the maximum competition contributed by a neighbor at the same position is 1, and the competition strength then linearly declines to 0 at the competition radius (default 0.01). Each fertile female has 20 times to choose any male to mate with in a “migration rate” radius (with a default of 0.04), and the females would not lay eggs if they haven’t found any male during this. Female fecundity is affected by competition strength in a 0.01 radius around the female (*ρ_i_*), a expected competition constant (*ρ*, proportional to population carrying capacity/total area, see below), and population growth rate in low density (*β*), which together determine female fecundity *ω_i_* = *β*/[(*β*-1) (*ρ_i_/ρ*)+1]. The number of offspring conform to a binomial distribution, with 50 independent chances to generate an offspring and a success probability for each equal to *ωi/25*. Offspring displacement from the mother are drawn from a normal distribution with a mean displacement of 0 and standard deviation equal to the migration rate. This produces an average displacement equal to *migration rate* * 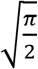. The position of offspring outside the boundaries is redrawn until they fall within the boundaries. We end the simulation 1000 generations after the drive release if the population or the drive were not eliminated earlier.

### Competition in the discrete-generation model

We introduce another species to represent competition between species with overlapping ecological niches. In this model, two species have the same reproduction rules, but the competition strength experienced by each individual is from not only the same species (intraspecies), but also from the other competing species (interspecies). However, because each species is somewhat specialized, competition from individuals from the other species will be reduced compared to competition from individuals of the same species. Hence, a new parameter called the competition factor (φ) is introduced into the model to govern the relationship between the two species. This multiplies interaction strength from the other species. In some scenarios, we also separate the interspecies competition factor for each species for cases when they might have asymmetrical competitive interactions. We release a gene drive only in the “target species” and refer to the other species as the “competing species. Accordingly, the expected competitions are as follows:

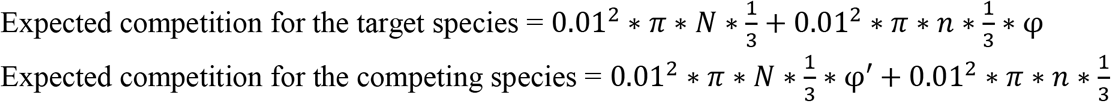

0.01 is the maximum competition radius (fixed for both species). *N* and *n* represent the total population carrying capacity for the target species and competing species, respectively. φ represents the interspecies competition factor that the target species experiences from the competing species, and φ’ that the competing species experiences from the target species. The factor of 1/3 comes from the linearly declining interaction strength between individual in the same position and individuals at the maximum competition radius.

### Mosquito model

We also used a model designed to simulate *Anopheles* populations in a previous study where it is discussed in more detail^20^, though we here consider it a more general mosquito model because most mosquito vectors have highly similar lifecycles. In the model, time steps are weekly, and individuals are considered to be at an immobile juvenile stage for the first two weeks. Instead of affecting female fecundity, competition here affects offspring survival. Juvenile mortality is determined based on competition from nearby juveniles after they are created, but not the next week. These individuals face the competition from other new offspring and week-old larvae nearby. These older larvae exert five-fold competition strength to individuals compared to new larvae because they consume more resources than new offspring. The expected competition between individuals is affected by the number of adult females in the population (10,000 by default) and the expected number of offspring per adult female per week (25, with females have a 50% chance to lay eggs each week, and each batch averaging 50 eggs), calculated as follows:

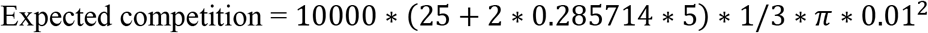

After calculating the actual competition experienced by a new offspring, we divide by the expected competition to obtain the competition ratio (*r*), which will determine the survival rate:

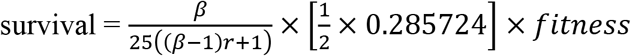

Where *β* is the low-density growth rate as before. In these equations 0.285724 is a constant that results in a fixed number of adult females at the specified value (10,000). Fitness is from direct fitness costs of the drive and does not include effects of somatic Cas9 expression.

Based on laboratory observations^28^ and to allow simulation of age-based health, the survival rates for adult mosquitoes are not density dependent, and female adults live longer than males. In our model, males live no longer than five weeks, and females never survive beyond their eighth. We use the following survival rates for adults advancing in age at each stage (the first two weeks represent juvenile stages):

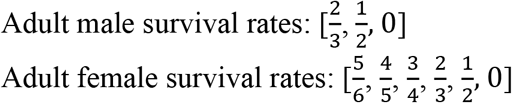

Adult females can attempt to mate from their first week, and they then immediately have a 50% probability to successfully produce offspring using the same sperm, as well as the same 50% probability in each subsequent week. We assumed female mosquitoes only have a 5% chance each week to re-mate since most mosquitoes usually mate only once in their lifetime^29–32^. Fertile females lay eggs based on Poisson distribution with the average of 50, which is multiplied by the fitness from leaky somatic Cas9 expression for drive individuals. At the end of each week, larvae experience density-dependent competition (see below), and all surviving adults have the same migration rate as offspring dispersing around their mother, which we set as 0.0307 by default (corresponding to the same per-generation allele advance rate as in the discrete generation model). Our mosquito density and migration is broadly consistent with previous field studies and models^20,33^ and perhaps represents an approximately 3.3 x 3.3 km area for *Anopheles gambiae*, though this would likely be reduced for other mosquitoes such as *Aedes aegypti* that have lower migration rates.

Simulations are run for 1,000 weeks after releasing the gene drive individuals.

### Competition in the mosquito model

The interspecies competition is essentially the same in the discrete generation and mosquito models. We assumed both species have the same life cycle and that competition only take place at the larval/juvenile stage. Therefore, the survival rate of both species is set in the same way as in the mosquito model, except that expected competition is now calculated as follows:

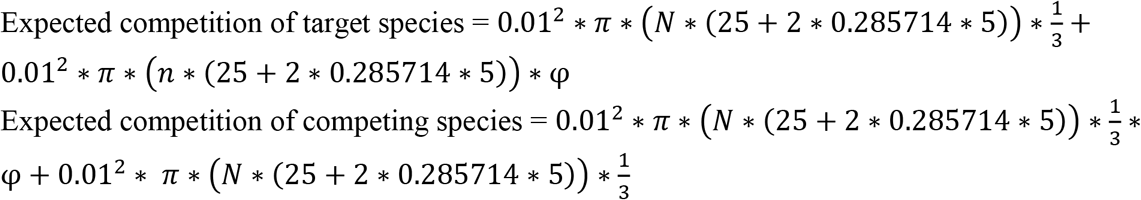

As before, *N* and *n* represent the number of adult females in target species and competing species, respectively. φ is the interspecies competition factor that the target species experiences from the competing species, and φ’ is the interspecies competition factor that the competing species experiences from the target species.

### Varying competition in different regions

In addition to a model where both competing species have the same characteristics throughout the arena, we developed a model where the space is evenly divided into two regions, with each species having an advantage in one region. Expected competition thus reverts to the same as in the one species model. For the discrete generation model, the equation is:

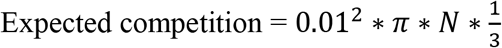

The competition factor φ still varies between 0 and 1 and determines the reduction in competition experienced by species in their preferred habitat from members of the other species. Interspecies competition from the species with the advantage remains at full strength, as does intraspecies competition) regardless of whether the species are in the region in which they have advantage or not).

We also tested a variant of the mosquito model where instead of varying competition, each female would have a reduced chance to reproduce if in the less preferred region of the species.

### Predator-prey model

Besides competition relationships, two species can also have a predatorprey relationship. We considered using gene drive to target either predator or prey separately in the discrete-generation model. Both species have the same reproduction rules in the discrete-generation model. The predators have the same effect on the prey species as a competing species, but with “predation intensity (θ)” replacing competition factor, which affects how deadly the predator is for the prey (we allowed this factor to go well above 1, which was the maximum for the interspecies competition factor). The equations are as follows:

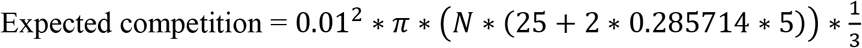

Compared to prey, predators have twice the migration rate and interaction distance, which affects interaction not only between predators, but also predators and prey. Similarly, predators have half the low-density growth rate that prey have (set to 5 as default). We set predator population size as 10,000 by default (one fifth the default size of the prey population). Predators experience normal competition with other predators, but the overall level of “competition” is adjusted by the amount of prey available. The extent to this occurs is governed by the “predator resource fraction” (λ), which represents how much of the predator’s diet is made up of the specific prey species we model. The specific equations are as follows:

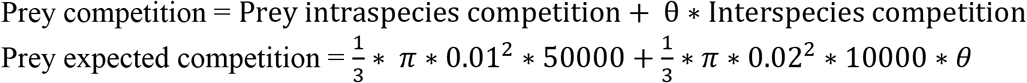

We also assumed the existence of predators in mosquito model (here, the target mosquito species can only be prey, not predators). Due to the wide variety of species that prey on all stages of mosquitoes, we assume for simplicity a generic predator that preys on mosquito larvae, where we already evaluate competition. The effect of predators on mosquito “competition” strength is as above (though the increased competition will affect juvenile mortality rather than adult fecundity as in the discrete-generation models). The existence of predators affects the expected competition, calculated as the equation:

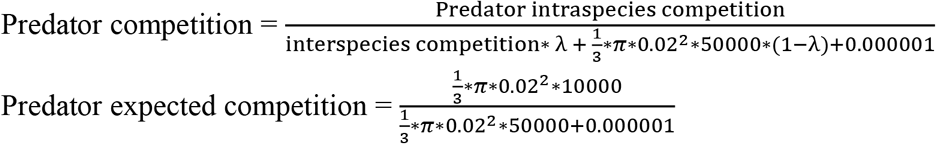

In our predator model, only 25% of female predators reproduce each week (half the rate of mosquitoes). To determine the number of offspring if they do reproduce, we use a binomial distribution as in the discrete-generation model but with 10 possible offspring. Adult mosquitoes and 1-week-old larva have normal interspecies interaction strength with predators, but 0-week-old larva only count for 1/20 as much strength (due to their smaller size). Predators are considered to be adults after their first week, and their survival rates are set so that their population linearly declines and so that they have twice the average generation time as mosquitoes. Additionally, no predators survive past their 12^th^ week. The expected competition of mosquito predators involves the number of predators (10,000 as default), as well as the number of prey adult females when the population is at capacity (also 10,000):

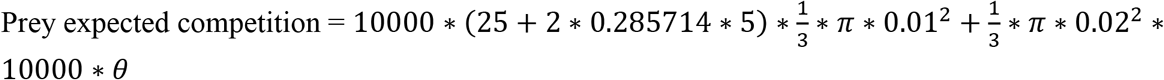

### Seasonality in the mosquito model

Mosquitoes often experience substantial seasonality, with high populations in the wet season and low populations in the dry season. To simulate this, we divided our weekly time steps into 52-week cycles. During these cycles, the population capacity of 10,000 fertile females was changed in all equations to a function with maximum and minimum determined by the seasonality constant:

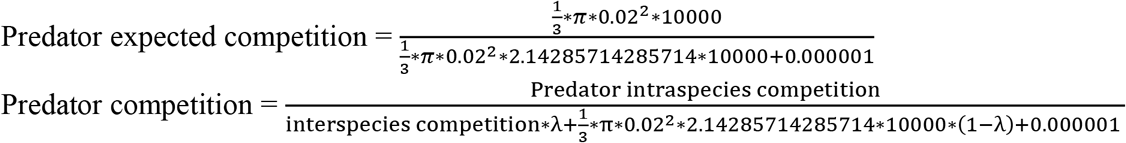

In one form, the population varied according to a sinusoidal function. In another form, the population capacity remained at its maximum or minimum levels for 22 weeks, followed by four weeks of linear transition to the other model. Drive individuals were released when the population capacity was at the baseline level of 10,000 and increasing.

### Assessment of the “chasing” phenomenon

To verify the degree of individual’s aggregation, we divided the 1×1 arena into 8×8 spots and counted the number of individuals in each spot to s^2^ calculate Green’s coefficient of each simulation. Green’s coefficient (*G*) is given by 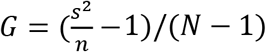, where *n* is the mean number of individuals around one grid, *s*^2^ is the variance of the counts and N represents the total population size. If individuals are placed randomly and according to a Poisson distribution, then it is expected that *n* = *s*^2^ and *G* = 0. Similarly, if all individuals highly aggregated into a cluster, then *G* =1.

When the drive is released in the center of the population, wild-type individuals are rapidly suppressed. Then, surviving wild type individuals are clustered near the boundaries, causing an increase in Green’s coefficient. Sometimes, suppression occurs at this point. However, previous results demonstrated that the suppression gene drive could result in “chasing” dynamics where the number of drive and wild type individuals fluctuates for a long time. This occurs when wild type individuals escape to an empty area and produce more offspring due to the lack of competition. Then, Green’s coefficient would decrease since these individuals occupy more area. We define chasing to start with the earlier generation of when Green’s coefficient first attains a local maximum value and when the relative number of wild-type individuals compared to the expected capacity first attains a local minimum (the comparison to expected capacity prevents seasonal fluctuations from potentially triggering this criteria).

### Data generation

The High-Performance Computing Platform of the Center for Life Science at Peking University was used to run the simulations. We used Python to process data and prepare the figures. All SLiM models and raw data are available on Github (https://github.com/jchamper/ChamperLab/tree/main/Suppression-Competition-Modeling).

## Results

### Drive characteristics and two-species models

We previously modeled highly efficient homing suppression drives in spatially continuous models, finding that with sufficient efficiency, successful and rapid suppression was usually possible^15,17,19,20^. In this study, we focus on a less efficient drive. Specifically, our default drive has a drive conversion efficiency of 80%, a 10% germline resistance allele formation rate, a 10% embryo resistance allele formation rate (caused by maternally deposited Cas9 and gRNA in the progeny of drive females), a 10% somatic fitness costs in females, and a 5% direct fitness cost in drive homozygotes (with multiplicative costs per allele). Such an imperfect suppression drive would be expected to reach a high equilibrium frequency in the population, potentially eliminating it if enough females are sterile at this point.

The same drive, in our discrete generation panmictic population model, would still be able to suppress the population (Figure S1). Despite having a genetic load of 0.85 (the fractional reduction in reproductive capacity compared to an equivalent wild-type population) at the drive’s high equilibrium frequency, this drive would not be able to suppress an infinite population with our default low-density growth rate of 10 (for which a genetic load of 0.9 would be necessary). However, stochastic effects would still allow the drive to successfully suppress the population in in all cases, though this often takes an extended period of time (Figure S1). In any given region of a spatial population, stochastic effects would be exaggerated, theoretically allowing suppression to occur even more quickly. Note that in our mosquito model, population suppression never occurs under these conditions (Figure S1), despite the same low density growth rate. These mosquito populations are more robust due to differences in the population dynamics due to the weekly time steps and viability-based competition. Specifically, the larger number of larva initially created tends to reduce stochasticity in this model compared to the discrete generation model.

In our previous discrete-generation spatial models, however, a drive with these default performance parameters would always fail to suppress the population and instead result in long-term “chasing” dynamics^15^. During chasing, the drive suppresses the population in a local area, but then the area is recolonized by a few wild-type individuals that escape the drive from adjacent areas. Due to low competition, these individuals can reproduce very quickly. The drive is still present and will reinvade the region, but complete suppression over the entire space is often never achieved. Even releases of better drives with a genetic load higher than 0.9 (sufficient to eliminate deterministic panmictic populations) often lead to this outcome, which is even more common in the mosquito model^20^, though sufficiently high drive efficiency will produce more successful outcomes^15,20^.

Nevertheless, drives with higher efficiencies (or at least lower fitness costs) may be difficult to construct. Successful gene drive-induced suppression may still be possible with our default drive, however, if a second species can also put pressure on the target species. Here, we examine three different two-species scenarios. In the first, two species coexist in the same region (Figure 1A). They have different ecological niches, and thus compete with each other to a limited degree. In the second scenario, we divide the arena into two regions, and each species is specialized to one of these regions (Figure 1B). Normally, each species would be mostly confined to their own region, with migration resulting in some limited overlap at equilibrium. In the third scenario, we model separate predator and prey species (Figure 1C), where the prey suffers in the presence of predators, and the predators require the presence of prey to reproduce at a normal rate.

**Figure 1.**
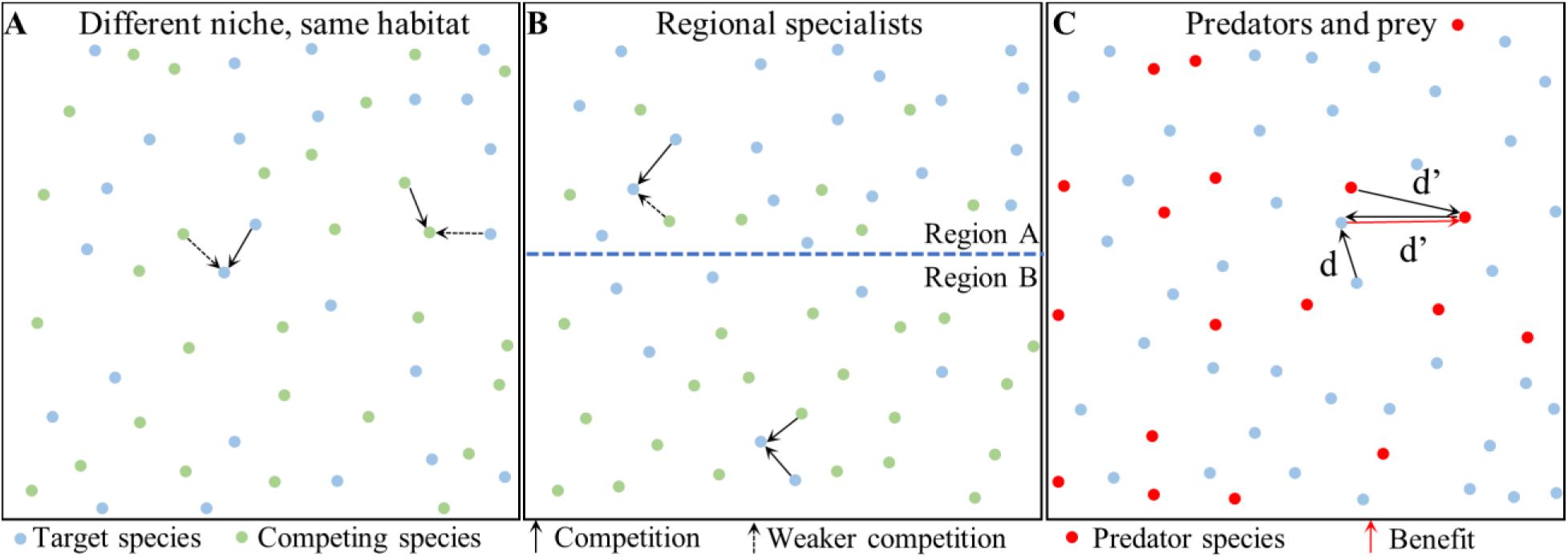
Two-species scenarios. (**A**) In the first scenario, two different species occupy different ecological niches in the same habitat. They have some overlap, so there is some interspecies competition, but intraspecies competition is greater. The competition factor determines the relative amount of interspecies competition. (**B**) In the regional specialist scenario, the arena is divided into two halves. In region A, all individuals experience normal competition from members of the target species and reduced from the competing species. In region B, this is reversed. The competition factor determines the relative amount of competition from individuals in their less preferred region. At equilibrium, the population of species A is substantially higher in region A than in region B. (**C**) In the predator prey model, prey individuals experience competition from both other prey and predators. Depending on the predation intensity, the competition from predators could be substantially higher. Predators experience normal competition from other predators. The presence of prey, however, promotes growth of predators, depending on the predator resource fraction (representing the importance of the prey species in the diet of the predators).

### Competing species promote population suppression

To assess the potential influence of a competing species on suppression drive outcomes (Figure 1A), we first implemented two different species in our discrete-generation model. The two species share the same habitat but occupy different ecological niches, so there will be some resource competition between them. In this model, we simplify the competition and life cycle characteristics of the two species, adjusting fecundity based on total competition from nearby individuals. Competition is determined by the number of individuals around the target species, which is separated into intraspecies competition and interspecies competition. Our model uses a competition factor variable to represent the relative interspecies competition strength between two species (0 = no competition, 1 = interspecies competition is as strong as intraspecies competition, an inherently unstable situation). When the target species is suppressed, the competing species tends to somewhat increase in population in the areas where the target species is no longer present. This can make it more difficult for chasing to occur, since the target species could still experience substantial competition in areas cleared by the drive that it recolonizes.

We first assumed that the competition factor is equal for both species (symmetrical competition), varying the population size of the competing species and the competition factor (Figure 2). Because our drive had modest efficiency, lack of competition invariably results in long-term chasing dynamics, with the drive still chasing after 1000 generations, similarly to our previous results^15^. With an increasing number of competing species and higher competition factor, it’s easier to suppress the target species, with the competition factor having a substantially larger effect in our parameter range. As the competition factor increases, there’s a small region of suppression after a period of chasing (Figure 2), though by the time the competition factor reaches about 0.5, the dominant outcome is usually suppression without chasing. For simulations where chasing occurred before suppression, the period of chasing was usually short, but when competing species size and competition factor are low, chasing could last for several hundred generations (Figure 2). Though chasing can make drives more vulnerable to complete failure by functional resistance allele formation^15,17^, the existence of this phenomenon does not prevent potentially substantial benefits from being achieved, depending on the specific drive application. Specifically, reductions in female reproductive capacity could reduce the fertile female population compared to its initial state (Figure 2). During short chases, patchy suppression tends to result in a higher number of average fertile females, but for more important longer chases, conditions that increase the drive’s efficiency or other factors that put pressure on the population such as high competition tend to reduce the average number of fertile females.

**Figure 2.**
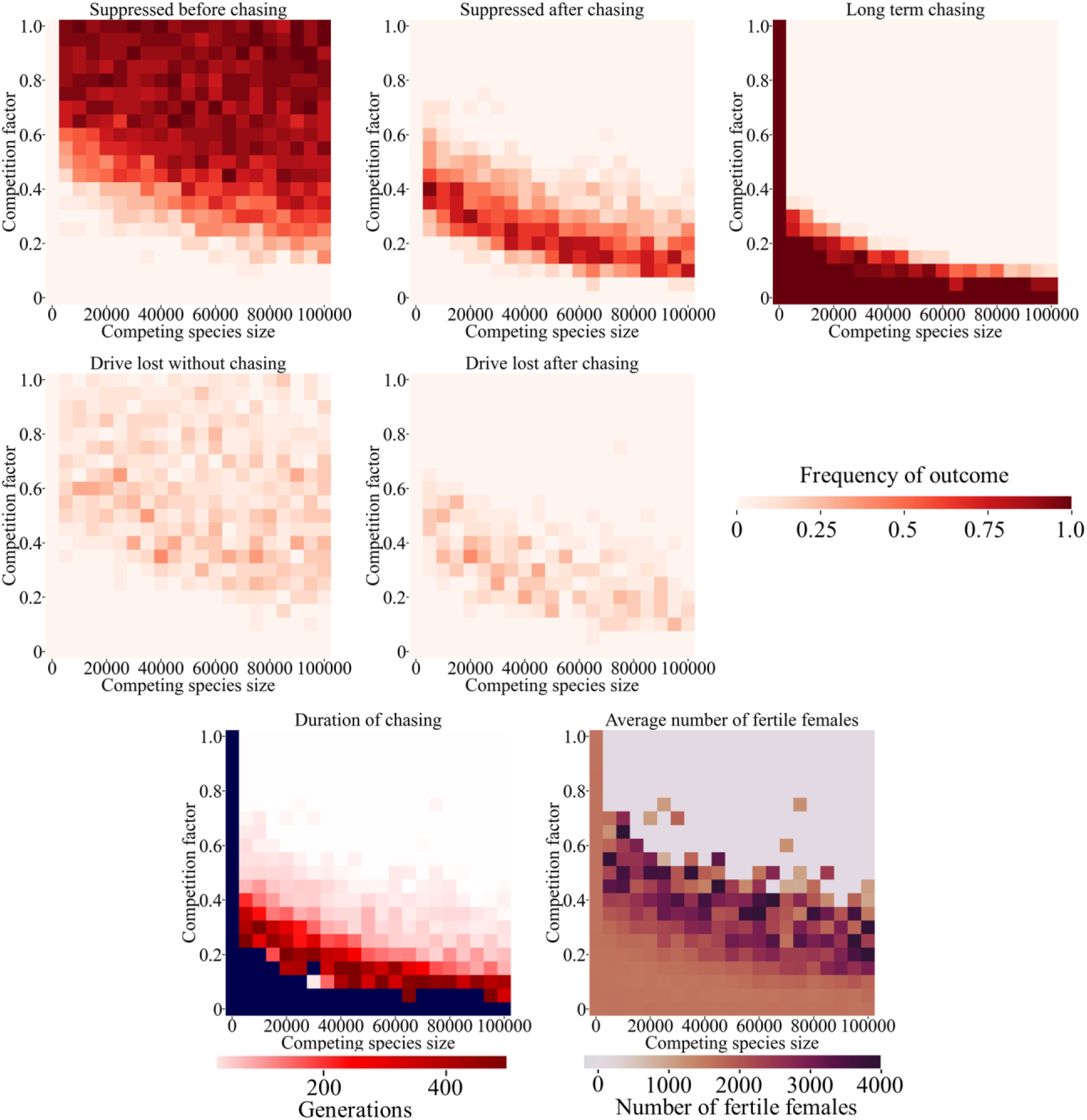
Competition in the discrete-generation model. Drive heterozygotes were released into the middle of a spatial population of 50,000 individuals of the target species that were colocalized with a competing species with varying population size and competition factor. The frequency of ultimate drive outcomes, the duration of chasing, and the average number of fertile females during chasing is displayed. Blue color refers to points where chasing continued to the end of the simulation in all cases. 20 simulations were assessed for each point.

Another mode of suppression drive failure involves stochastic loss of the drive allele. This is because gene drive individuals may struggle to find a potential mate in a limited distance after the population is sufficiently suppressed, removing the gene drive form the population while clusters of wild-type individuals still remain (at which point they could quickly repopulate the arena). This outcome is usually rare in one-species models for suppression drives targeting female fertility genes^15,17^. However, drive loss is substantially more common in our model of competing species (Figure 2). Though never the dominant outcome, it represents a substantial fraction for the entire parameter range except when competition between the species is very low (when long-term chasing outcomes dominate).

Another potentially important scenario is asymmetric competition, where species affect each other differently. To assess this phenomenon, we adjusted the competition factor separately for the target and competing species. As expected, we found that competition from the competing species on the target species (competition factor B) was helpful for suppressing the target species (Figure 3). As before, a modest increase in this competition factor above zero was sufficient to prevent long-term chasing, and further increases could reduce the frequency and duration of shorter-term chasing (Figure 3, S2). Drive loss remained a small but significant outcome over most of the parameter range.

**Figure 3.**
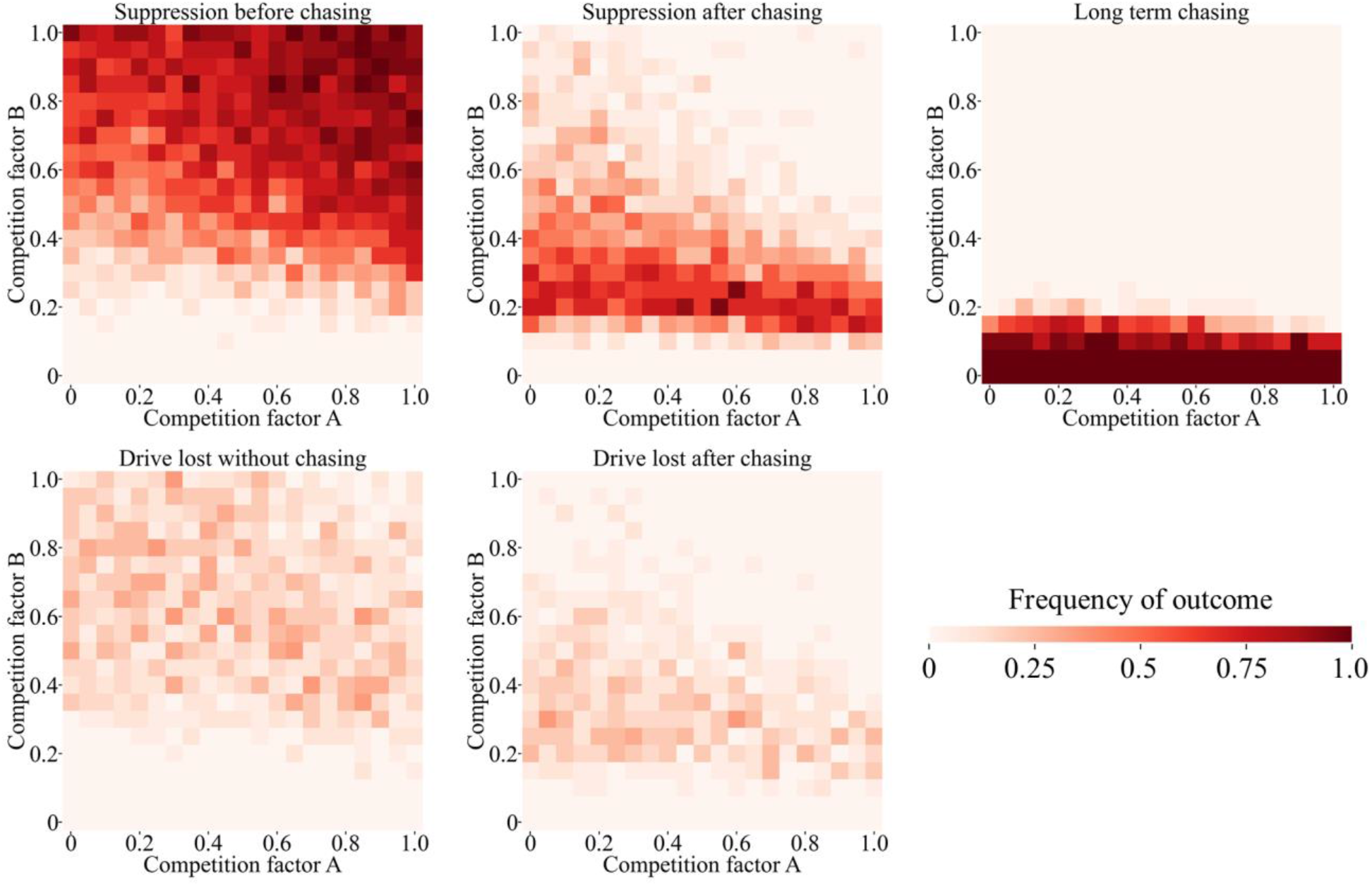
Asymmetric competition. Drive heterozygotes were released into the middle of a population of 50,000 individuals of the target species that were colocalized with a competing species with 50,000 population size. The relative degree of competition between the species was varied, with competition factor A referring to the competition experienced by the competing species from the target species, and vice versa for competition factor B. The frequency of ultimate outcomes is displayed for each simulation, and 20 simulations were assessed for each point in the parameter space.

Surprisingly, we found that the competition factor A (the competition from the target species experienced by the competing species) had little effect in the range tested (Figure 3, S2). Potentially, higher competition would allow wild-type members of the target species to better push back against the competing species after temporary suppression to reclaim its higher population. This could facilitate chasing. However, we found that higher competition factor A tended to slightly reduce instances of suppression after chasing in favor of suppression without chasing. This may be because it enabled gene drive individuals to push back for slightly longer against encroaching members of the competing species, giving them more time to mate with adjacent wild-type individuals to spread the drive.

We previously developed a mosquito-specific model, which indicated that successful suppression would be substantially more difficult than in our discrete-generation model^20^. Because most mosquito vectors have broadly similar lifecycles, we used a simplified model in which the two species have the same reproduction rules, and we assumed symmetric competition. As in the discrete-generation competition model, suppression without chasing was the most common outcome when competition factor and competing species population was high, shifting to suppression after chasing and eventually long-term chasing (Figure 4). However, compared to the discrete-generation model, long-term chasing occupied a much greater fraction of the parameter range. Furthermore, drive loss become a major outcome, more common than suppression after chasing in the intermediate parameter range. This is unexpected because in previous studies with one species, drive loss was less common in the mosquito model than in the discrete-generation model^20^. When the competition factor was low, chasing could last for a long time with a high average population of adult fertile females, despite the lower initial population compared to the discrete-generation model (Figure S3).

**Figure 4.**
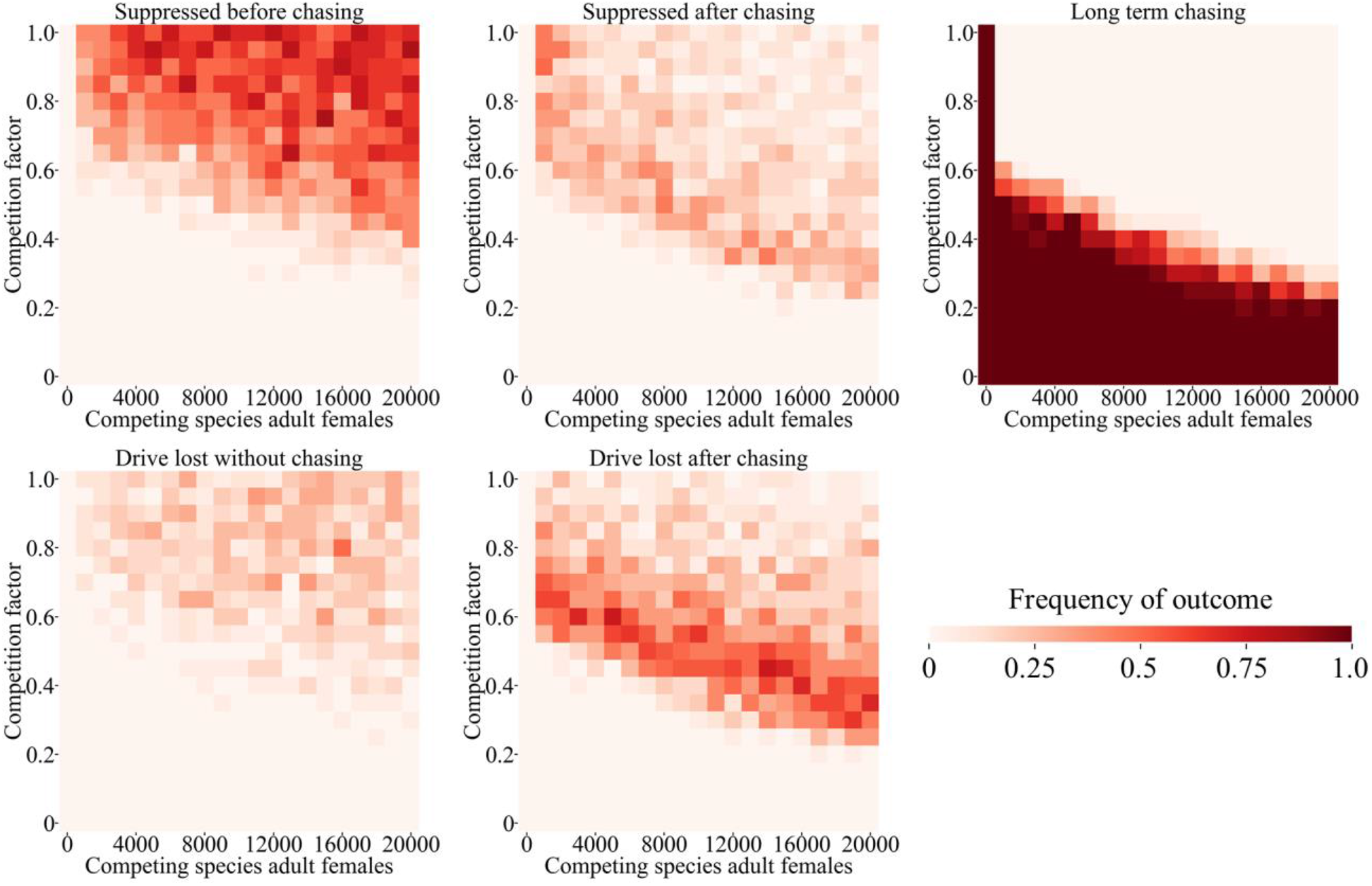
Competition in the mosquito model. Drive heterozygotes were released into the middle of a mosquito population with 10,000 adult females of the target species that were colocalized with a competing species with varying population size and competition factor. The frequency of ultimate outcomes is displayed for each simulation, and 20 simulations were assessed for each point in the parameter space.

### Regional specialists

For many competing species, instead of being specialized to particular ecological niches within a region, they are specialized to certain regions themselves in a variety of ways. We modeled this situation by dividing our arena into two equal halves, representing different environments (Figure 1B). Each of the two competing species are specialized to one of these environments. In their preferred environment, the species compete normally, but their competition factor is reduced for interspecies competition when in their non-preferred environment. In this scenario, the migration rate is also important because it will determine the initial degree of population overlap between the two species, together with the competition factor. As expected, higher migration results in more suppression outcomes, while low migration results in more drive loss outcomes (Figure S4), consistent with previous results^15,17,20^. Competition also had similar effects in this environment, with a modest level of competition allowing suppression without chasing in most cases or shorter chases, as long as migration was not very low.

In our mosquito model of two regional specialists, we found that the general pattern compared to the discrete-generation model was mostly the same (Figure 5, S5), though higher levels of competition and migration were required to avoid long-term chasing results. Similar results were seen in scenarios when larvae had an advantage in their preferred region (Figure 5) and when females had a higher chance of reproduction in their preferred region (Figure S6). This latter phenomenon could be caused by different mosquito species being specialized to target humans or other animals and thus having different reproductive success in natural environments and those with high human populations.

**Figure 5.**
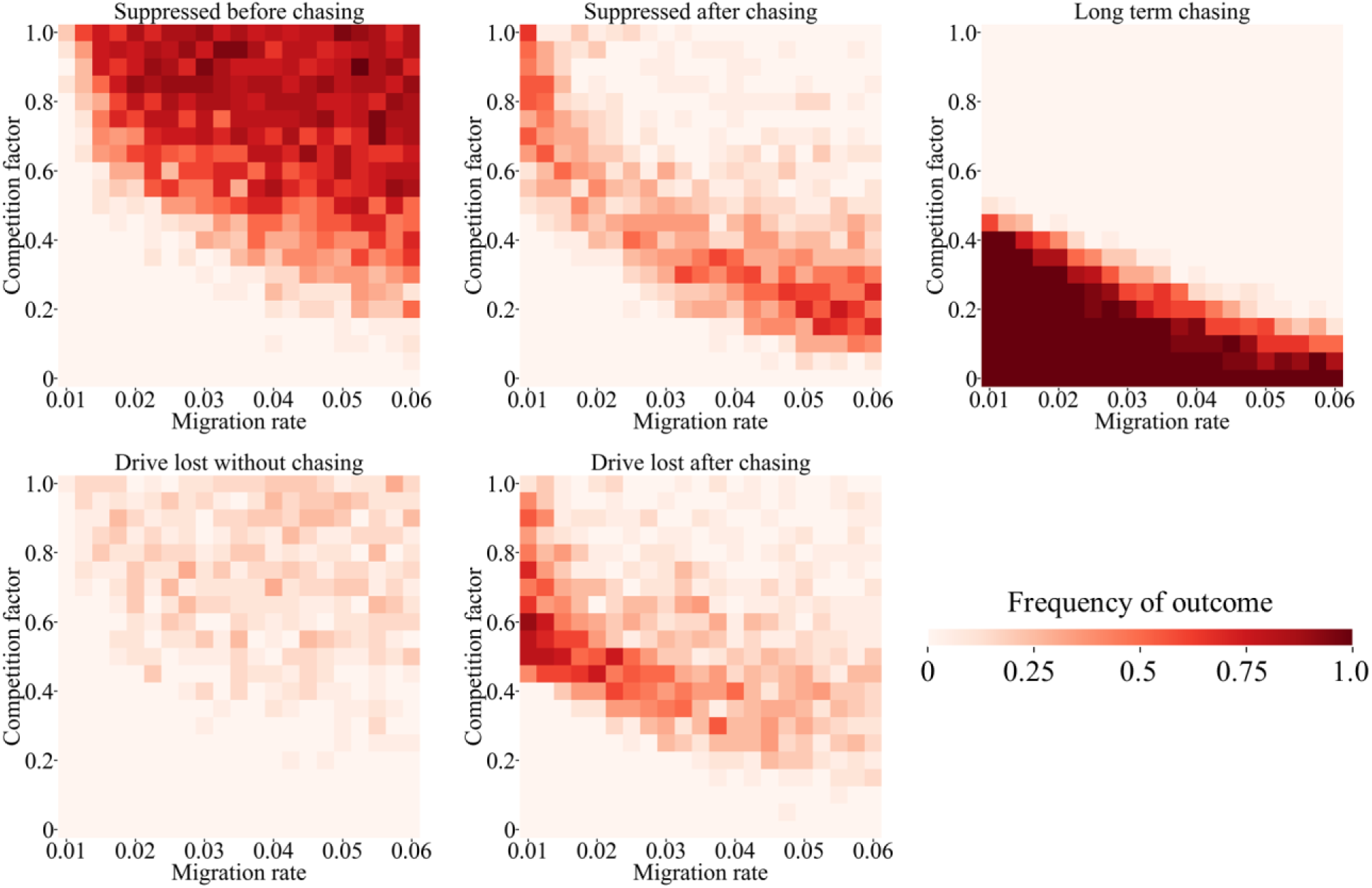
Two-region mosquito model. Drive heterozygotes were released into the middle of a space divided into two regions with two mosquito species with varying migration rate and competition factor. Each species is specialized in one region and competes normally there, while competing at a reduced rate determined by the competition factor in the other region. The frequency of ultimate outcomes is displayed for each simulation, and 20 simulations were assessed for each point in the parameter space.

### Predator-prey model

Predator-prey relationships between species are also common, and this asymmetric scenario could be expected to affect gene drive outcomes differently depending on whether the predator or the prey is targeted. We modeled predator-prey relationships by assuming that predators provide competition to prey depending on the predation intensity, which could allow a predator to provide more “competition” to a prey than a member of the same species. Predators compete with each other and are also influenced by the predator resource fraction (representing the importance of the prey in the predators’ diet). Predators received a reproductive penalty when few prey individuals were nearby and a bonus if prey were plentiful.

As expected, high predation intensity values had a similar effect as a competing species (Figure S7), with even modest values being sufficient to allow for suppression without chasing (though still with some risk of drive loss). However, when the predator resource fraction was high, the population of predators tended to decrease after the suppression drive reduced the prey population. This could substantially reduce predation levels of the prey, allowing them to more effectively avoid the suppression drive in these cases. Even with high predation intensity, high predator resource fractions could effectively prevent suppression.

A gene drive could also be used to target a predator. In this case, the presence of the predatorprey system does not benefit the drive. Instead, suppression of predators could allow for an increase in the prey population (Figure S8), benefiting remaining predators. Thus, when assessing this, we allowed the drive conversion efficiency to vary, allowing for demonstration of the effect of predator-targeting on suppression performance. Fixing the predation intensity at 1, We found that at higher drive conversion, suppression outcomes became more common (Figure S8), as expected^15^. The predator resource fraction had little effect on these outcomes, only slightly reducing the chance of successful suppression when high (Figure S8). This is because the predators can benefit from slightly increased prey populations. Higher predation intensity could increase the magnitude of this effect.

Mosquitoes are often prey of many species, so to create a simple model, we implemented a generic mosquito predator that preys on the common larval stage. The predation intensity here has to be scaled upward compared to the discrete-generation for the predators to provide roughly the same relative competition compared to intraspecies competition (for example, a predation intensity of 1.25 means that predators provide as much average competition as intraspecies competition in the discrete generation model, but the corresponding predation intensity for larvae in the mosquito model is 6.96). Overall, we find a broadly similar pattern to the panmictic model (Figure 6), though with substantially higher predation intensity required for successful suppression to be possible, and even then, drive loss outcomes are often more common. Even moderately high predator resource fractions can fix the outcome at long-term chasing when predation intensity is high. However, while predators can certainly impact mosquito populations^34,35^, it remains unclear if any target mosquito species actually represents a substantial portion of the diet of any predator^35–37^, so lower values of this variable are likely more realistic.

**Figure 6.**
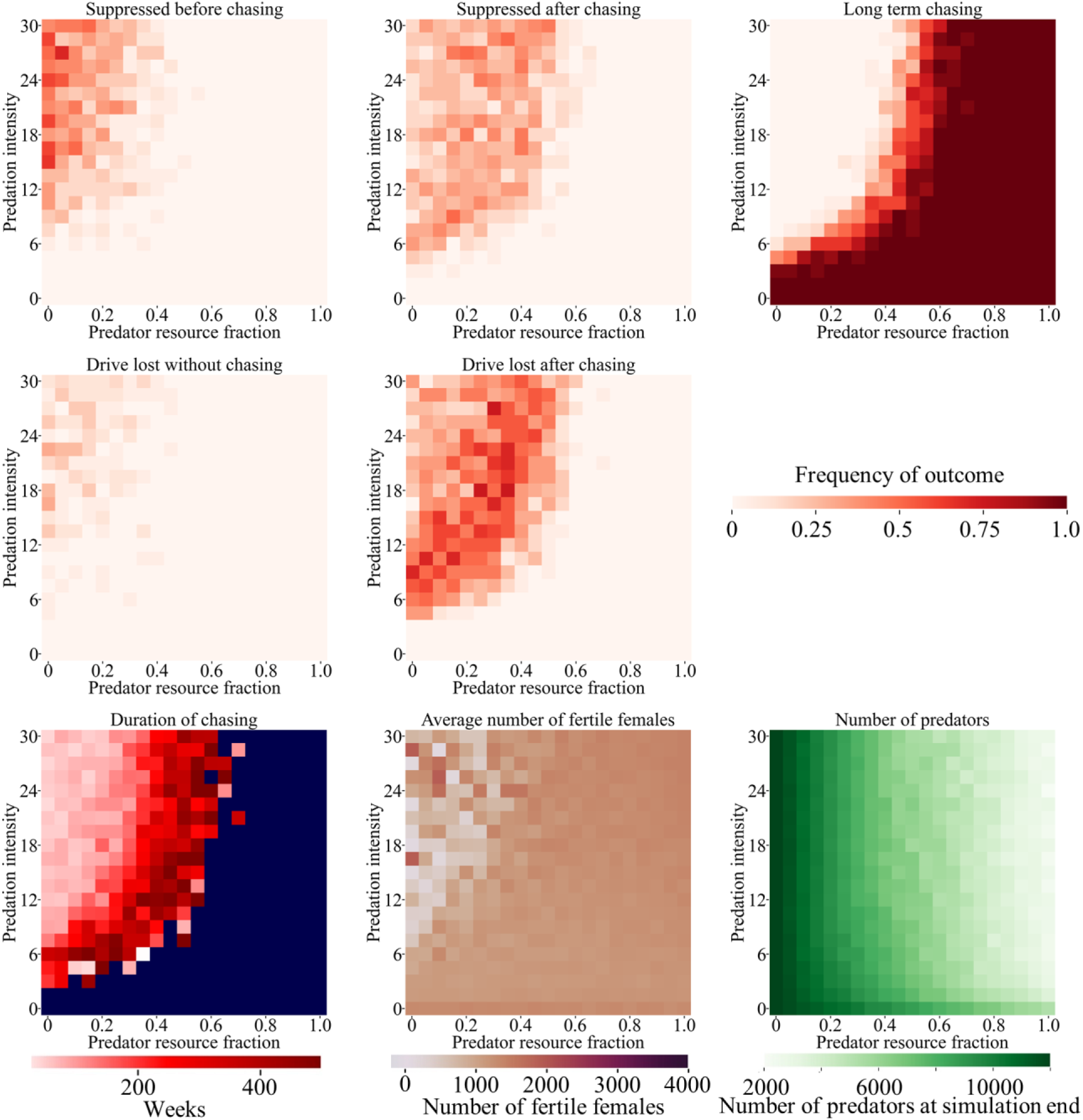
Predator-prey in the mosquito model. Drive heterozygotes were released into the middle of a mosquito population with 10,000 adult females of the target species that were colocalized with a predator population of 10,000 with varying predation intensity and predator resource fraction (representing the fraction of the importance of the target species to the predator’s diet). The frequency of ultimate outcomes, duration of chasing, average number of fertile females during chasing, and number of predators at the end of the simulation is displayed. Blue color refers to points where chasing continued to the end of the simulation in all cases. 20 simulations were assessed for each point in the parameter space.

### Effect of seasonality on the mosquito model

In many environments, mosquitoes (particularly *Anopheles* malaria mosquitoes) experience major population fluctuations based on the climate^26,38–41^. During the wet season, populations are high. Such populations can be greatly reduced or even locally eliminated during the dry season. These fluctuations could add a degree of stochasticity, potentially aiding gene drive-based suppression. However, at the beginning of a wet season, the pressure from competing species could be removed, allowing the target species to achieve high growth rates, which may increase the frequency of chasing.

To investigate this, we first incorporated seasonality into our single-species mosquito model, though we did not model extreme seasonality because this requires factors beyond the scope of our model to ensure mosquito persistence (such as long-distance migration or aestivation). The seasonality factor controls the magnitude of the population fluctuations, representing a proportionate increase during the peak of the wet season and a decrease during the low point of the dry season. Both sinusoidal models of population capacity fluctuation as well as sharp transitions between maximum and minimum population levels were considered. We found that successful suppression could only occur when drive conversion efficiency was high, and even then, long-term chasing outcomes remained possible (Figure S9–10). High population fluctuations due to seasonality of up to 80% did slightly increase the rate of suppression, and chasing could be avoided in slightly more instances with sharp transitions between seasons. However, higher seasonality also increased the average number of fertile females during periods of chasing.

In a competing species model, we set the population of the competing species as equal to the target species while allowing the competition factor and seasonality intensity to vary. In this situation, seasonality could allow the target species to be partly released from competition during the beginning of the wet season. Unlike the single-species model, increased seasonality intensity was usually harmful to the drive over most of its parameter range. When competition was high enough to potentially enable suppression, higher seasonality with sharp seasonal transitions reduced the rate of suppression without chasing, with these results generally converted into suppression after chasing (Figure 7). Drive loss outcomes were also similarly changed. However, these periods of chasing tended to be short (albeit with a high average number of fertile females), so the overall effect of seasonality was not substantial. These effects were also seen to an even lesser degree with the sinusoidal seasonal model with softer seasonal transitions (Figure S11).

**Figure 7.**
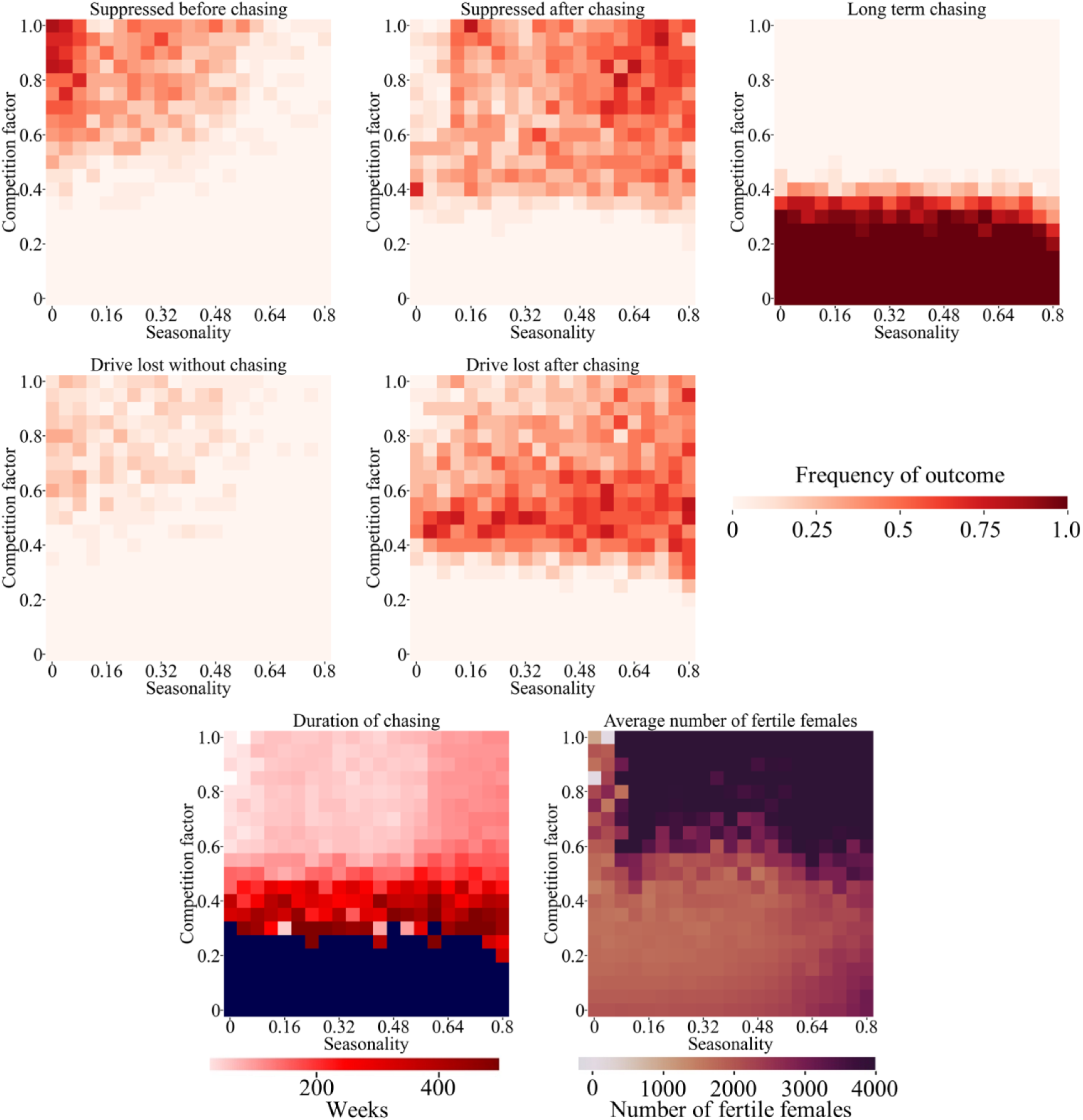
Seasonality and competition. Drive heterozygotes were released into the middle of a mosquito population with a baseline 10,000 adult females of the target species that were colocalized with a competing species with equal population size and a competition factor of 0.5. The population goes through a yearly cycle with the maximum and minimum being above and below the baseline by 10,000 x seasonality constant and a sharp, linear transition between these lasting for four weeks. The frequency of ultimate drive outcomes, the duration of chasing, and the average number of fertile females during chasing is displayed. Blue color refers to points where chasing continued to the end of the simulation in all cases. 20 simulations were assessed for each point in the parameter space.

### Simplified modeling two species

In this study, the main general effect of competing species or predators is to provide a constant level of competition, preventing the target species from achieving its potential low-density growth rate that would be possible in an ideal environment where only the target species is present. Though two-species interactions are often more complex, we can potentially construct a simple model of competing species by using only a single species with a reduced low-density growth rate. The calculation of this rate assumes complete absence of the target species from a region and potential growth of the competing species due to reduced competition from the target species at equilibrium. The adjusted low-density growth rate of the target species could then be simply calculated and set as the new low-density growth rate of the model. Such a model would not capture small-scale dynamics involving the process of replacement of the target species by the competing species, or resurgence of the target species in local areas against the competing species, but it could still approximate some interaction dynamics.

To assess how accurate such a simplified model would be, we first allowed the low-density growth rate to vary in the discrete-generation model (Figure S12). Because this drive had lower efficiency than our default one in our previous study^15^, we observed a more dramatic transition between long-term chasing and suppression without chasing as the low-density growth rate decreased. We then held the low-density growth rate at 10 in two sets of competing species simulations with competing populations of 50,000 and 5,000. In these simulations, the competition factor was adjusted to produce the same “equivalent” low-density growth rates when the competing species fully replaced the target species. We found that the outcomes of these two models were nearly identical, with the only major difference being a slight shift in the 5,000-population competing species data set from long-term chasing to suppression after chasing. This difference from the 50,000-population competing species model was likely due to increased stochasticity from the smaller competing population, which supported suppression. Thus, it may be possible to accurately simulate the outcome of suppression gene drive with certain two-species interactions by adjusting the low-density growth rate in single-species models.

## Discussion

Many spatially explicit models of suppression gene drive have showed that achieving population suppression may be considerably more difficult than expected compared to simpler panmictic models^15,17–19,26,42–44^. Population elimination in these more complex settings often requires gene drives with very high efficiency and low fitness costs, which has perhaps not been achieved in even the best *Anopheles* drives thus far^20^. However, in this study, gene drive performance was improved after we added additional complexity in the form or competing species or predators. We found that these factors could allow even a moderately efficient suppression drive to still successfully eliminate a target population. This suggests that population suppression by gene drive could be potentially more viable and efficient than was previously thought.

For long-term stability, each species must have its own ecological niche, which could take a variety of forms. However, it is rare for this niche to have no overlap with any other species, considering the wide variety of generalist and specialist species present in most ecosystems. When competing species or predators are present, a target species will not be able to grow at their theoretical maximum rate, even if intraspecies competition is low and resources are plentiful. Though these pressures alone are unable to cause extinctions under normal circumstances, the additional pressure of even a weak suppression gene drive may be enough to induce population collapse. In our discrete-generation model, higher growth rates are only achieved in the near absence of competition. Thus, gene drive-induced population suppression was successful even if the interspecies competition factor or predation intensity was low. In the mosquito model, the population is more robust, and achieving population elimination is more difficult^20^. Nevertheless, moderate levels of competition in this model could still ensure the success of a suppression drive, even one much weaker than those previously considered for *Anopheles*^20^. These results were robust to additional of seasonality, an important characteristic of mosquito population dynamics in many regions. This boost to suppression may be critical because high efficiency suppression drives have only been constructed in *A. gambiae*, and even these suffer from moderate to high fitness costs due to undesired somatic Cas9 expression. In mice^21,22^, *Aedes*^23,24^ and even *Drosophila*^6^, homing drives thus far have had lower drive conversion efficiency, likely too low for elimination of a robust population lacking competitors.

Our study also identified situations in which suppression may become more difficult. These include situations where a target prey species is exclusively preyed on by a predator (high “predator resource fraction). In this case, the predator may go extinct before the prey, releasing the prey from predation pressure and substantially reducing the likelihood of successful suppression. Attempts to suppress a predator may also be somewhat more challenging if partial removal of the predator releases the prey from pressure, increasing their population and providing more resources for the remaining predators. Finally, our results indicate that even though competitor or predation pressure increases the chance of successful suppression, they also increase the chance of drive loss. This is not necessarily a disastrous result for a suppression drive if drive can migrate from other adjoining regions where it is still present or if a small additional drive release is feasible. However, it could certainly make suppression more difficult and complex, particularly in more realistic environments, where drive loss outcomes may further increase in frequency if the population includes some semi-isolated patches that are more difficult for the drive to invade.

Two other undesired results are also suggested by our study. First, as we saw in our predatorprey models, extinction of the predator may become more likely if the predator specializes on the target prey. This outcome may be undesirable and should be considered when evaluating populations for suppression by gene drive or other methods. So far, an analysis of this for malaria mosquitoes has indicates that most predators would not be substantially affected by their removal^36^. Another potentially more serious issue would be a competing species increasing their population in response to a successful suppression strategy. In some cases, this would cause no problems (particularly if the competing species was a native species under threat by the invasive target species). However, for mosquitoes, the competing species might also be a major disease vector or have the potential to become one when occupying a new region. In this case, the benefits of suppressing only the target species might be substantially reduced compared to expectations. It might be necessary to eliminate several species in a region to accomplish the desired aim, which suggests that a targeted modification drive approach may be a simpler strategy if it were also feasible. It remains unclear to what extent this may be an issue for various mosquito species. For *Anopheles*, some vectors have already partially replaced *A. gambiae* after deployment of insecticide-treated bednets^36^. *Aedes albopictus* is also known to be present where *Aedes aegypti* is found^45^ and can also be a major disease vector, albeit a less efficient one. Another study found limited interactions between *Culex* and *Anopheles*^35^, though even in a situation where one species dominates and another avoids competition, the less adapted species could still replace an empty niche and somewhat inhibit the first species if it later returns to the niche during a period of chasing.

Though our study has further increased model complexity compared to previous work, it still lacks several factors that may prove important to the outcome of a suppression drive. Our discrete-generation model, of course, is a simple and non-specific representation of a spatially distributed population, and the differences in outcomes between it and our mosquito model already demonstrate the importance of factors such as lifecycle and type of competition. Our mosquito model has greater detail, but still involves a uniform landscape, random movement, and other approximations which may not be representative of actual populations, such as our simplified version of seasonal population fluctuations. In particular, any competing species may have different lifecycle and survival characteristics. Competition could take a different form as well, rather than larval competition or variance in adult bite rates by region. For example, mating between *A. aegypti* and *A. albopictus* may asymmetrically produce female sterility^45^. The geographic pattern of species advantages would also be considerably more complex than our division of the arena into two equal portions, and in many cases would involve smoother gradients of transition and patchy preferred environments. Predators can also be highly diverse and eat adults as well as larva, in addition to competing among themselves in different ways than we modeled. These factors can all potentially be investigated in future studies, though in many cases, more specific ecological information may need to be collected before accurate predictions can be made.

We have shown that multi-species modeling may be necessary for accurately predicting the outcome of a gene drive. However, this also substantially increases the complexity and run-time of the model. In many cases, it could be desirable to avoid such modeling while still incorporating the effects of different species. We showed that by adjusting the low-density growth rate (often called the “intrinsic growth rate”) in our model, we can get approximately similar outcomes to situations with multiple competing species. This does require some knowledge about the nature of interspecies interactions, and to retain accuracy in a model incorporating seasonality, additional adjustments would need to be made for the model to still accurately represent population dynamics. For example, the low-density growth rate could be allowed to go to its maximum at the beginning of a wet season and then decline afterward as maximum growth rates temporarily become impossible due to competition and/or predation.

We have shown that in many cases, modeling of competing species and predators may be necessary to accurately predict the outcome of a suppression gene drive release. Specifically, we found that these factors can perhaps make suppression substantially easier than anticipated by simpler models. While simple models are generally preferred when possible, increasing the detail and realism of suppression gene drive models thus far has continued to produce substantially different results, highlighting the complexity of this dynamic system.

## Acknowledgements

Thanks to the High-Performance Computing Platform of the Center for Life Science at Peking University for assistance with cluster-based data collection. This study was supported by laboratory startup funds from Peking University and the SLS-Qidong Innovation Fund.

## Supplemental Information

**Figure S1.**
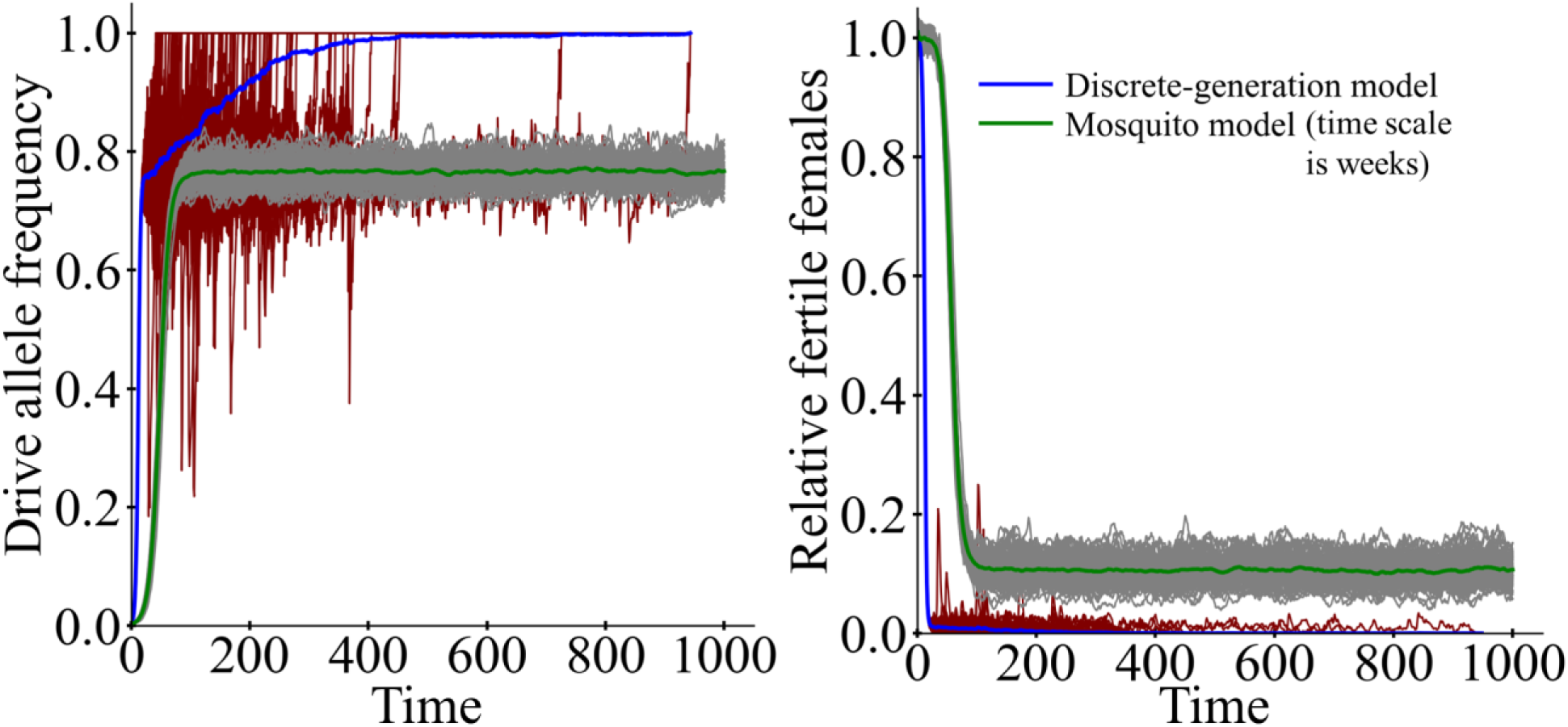
Performance of the moderate efficiency suppression drive in panmictic populations. Drive heterozygotes were released into a panmictic population of 50,000 individuals (discrete-generation model) or 10,000 adult females (mosquito model) of the target species. The drive allele frequencies, total population sizes, and relative fraction of fertile females (compared to the starting populations) are tracked. Thin lines represent individual simulations (100 total for each model), and thicker lines represent average values.

**Figure S2.**
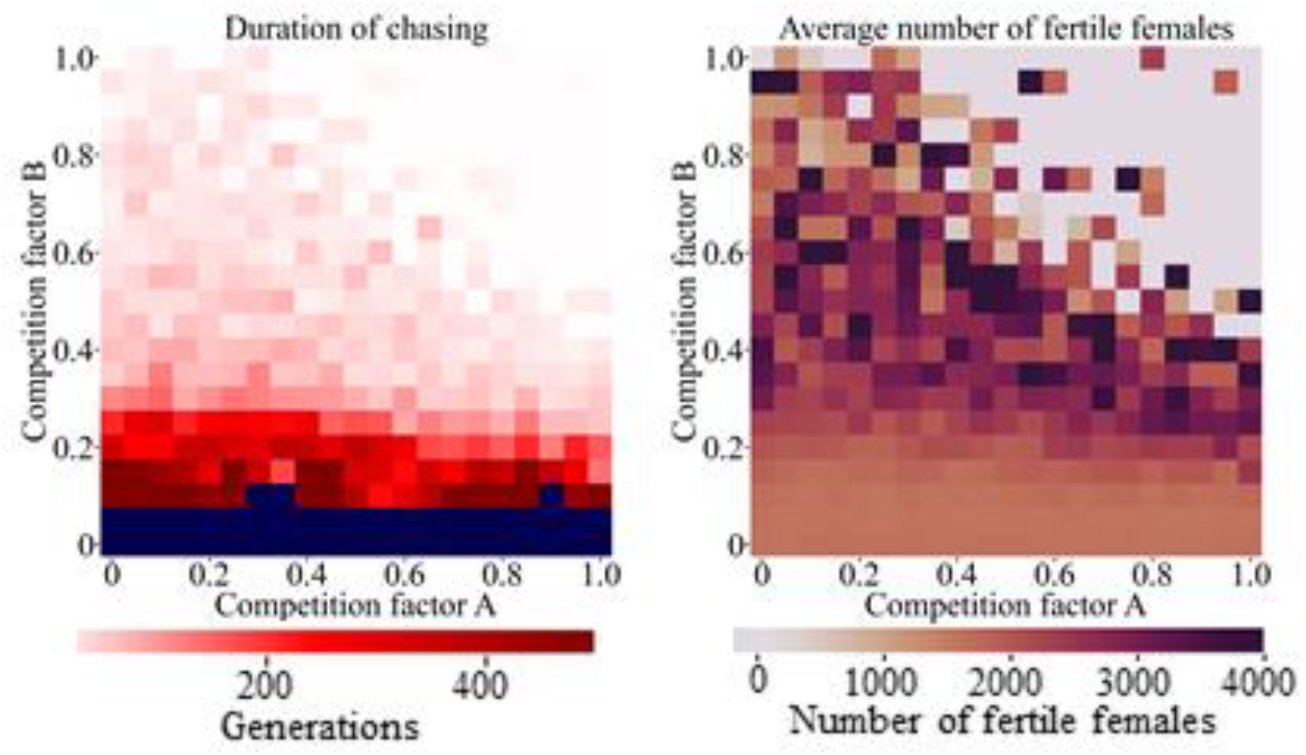
Asymmetrical competition. Drive heterozygotes were released into the middle of a population of 50,000 individuals of the target species that were colocalized with a competing species with 50,000 population size. The relative degree of competition between the species was varied, with competition factor A referring to the competition experienced by the competing species from the target species, and vice versa for competition factor B. The duration of chasing and average number of fertile females during chasing is displayed. Blue color refers to points where chasing continued to the end of the simulation in all cases. 20 simulations were assessed for each point in the parameter space.

**Figure S3.**
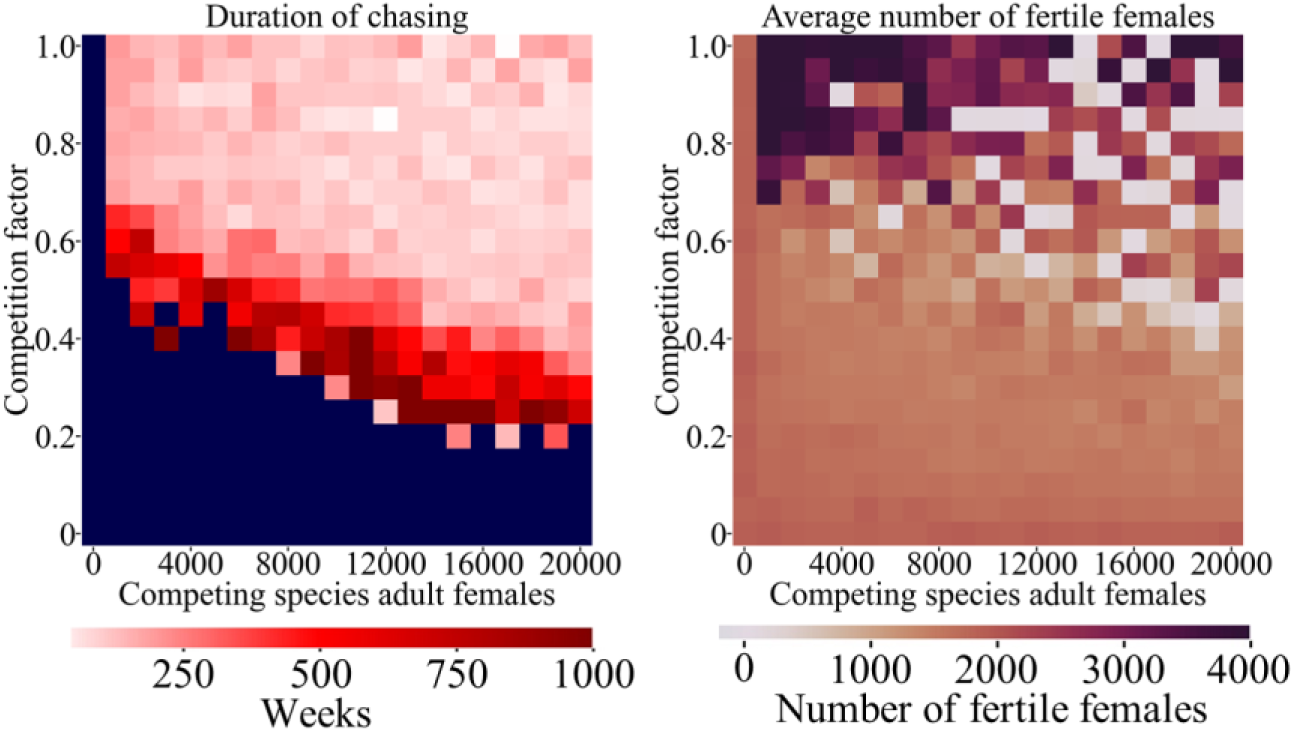
Competition in the mosquito model. Drive heterozygotes were released into the middle of a mosquito population with 10,000 adult females of the target species that were colocalized with a competing species with varying population size and competition factor. The duration of chasing and average number of fertile females during chasing is displayed. Blue color refers to points where chasing continued to the end of the simulation in all cases. 20 simulations were assessed for each point in the parameter space.

**Figure S4.**
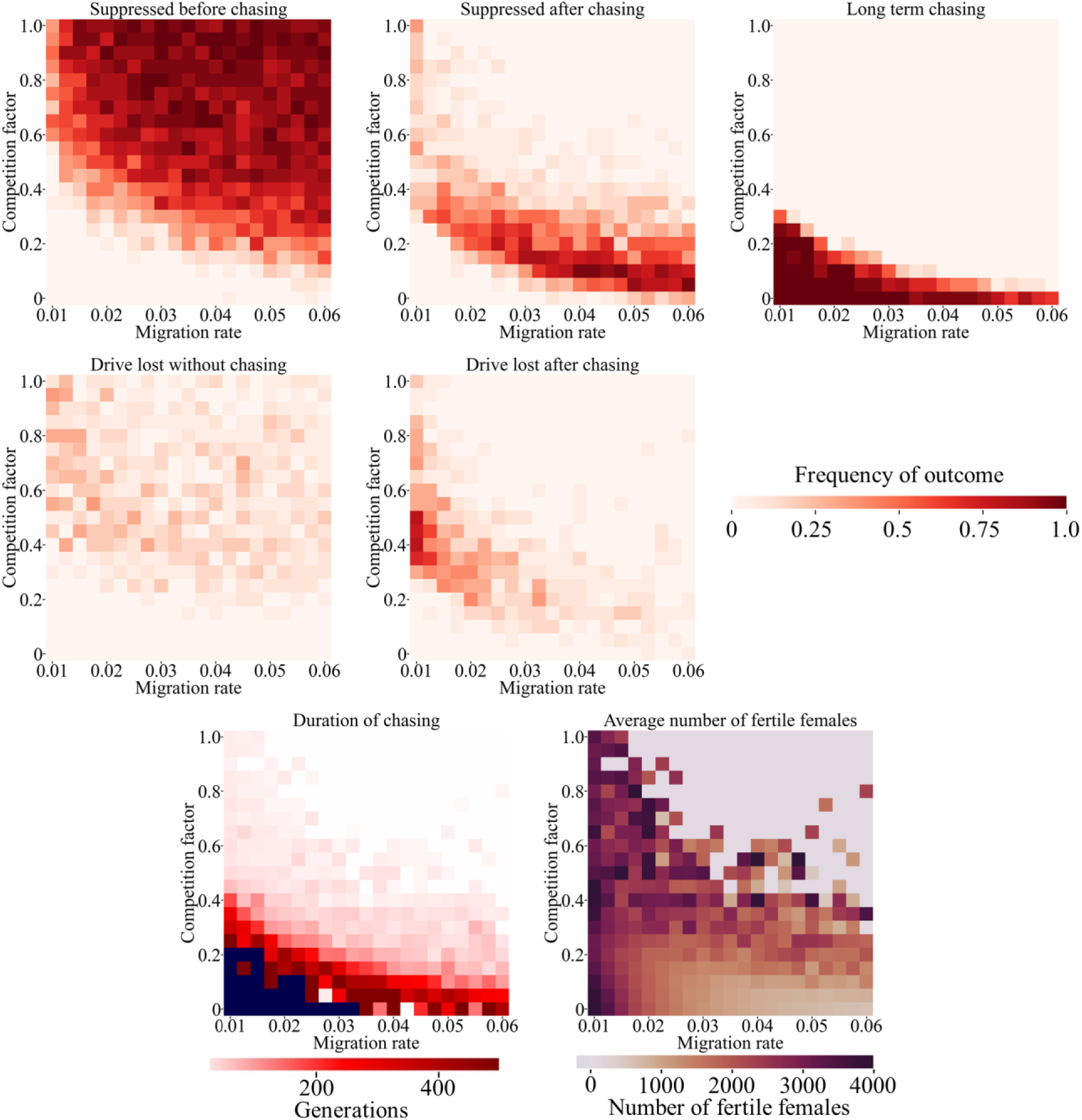
Two-region discrete generation model. Drive heterozygotes were released into the middle of a space divided into two regions with two species with varying migration rate and competition factor. Each species is specialized in one region and competes normally there, while competing at a reduced rate determined by the competition factor in the other region. The frequency of ultimate outcomes, duration of chasing and average number of fertile females during chasing is displayed for each simulation. Blue color refers to points where chasing continued to the end of the simulation in all cases. 20 simulations were assessed for each point in the parameter space.

**Figure S5.**
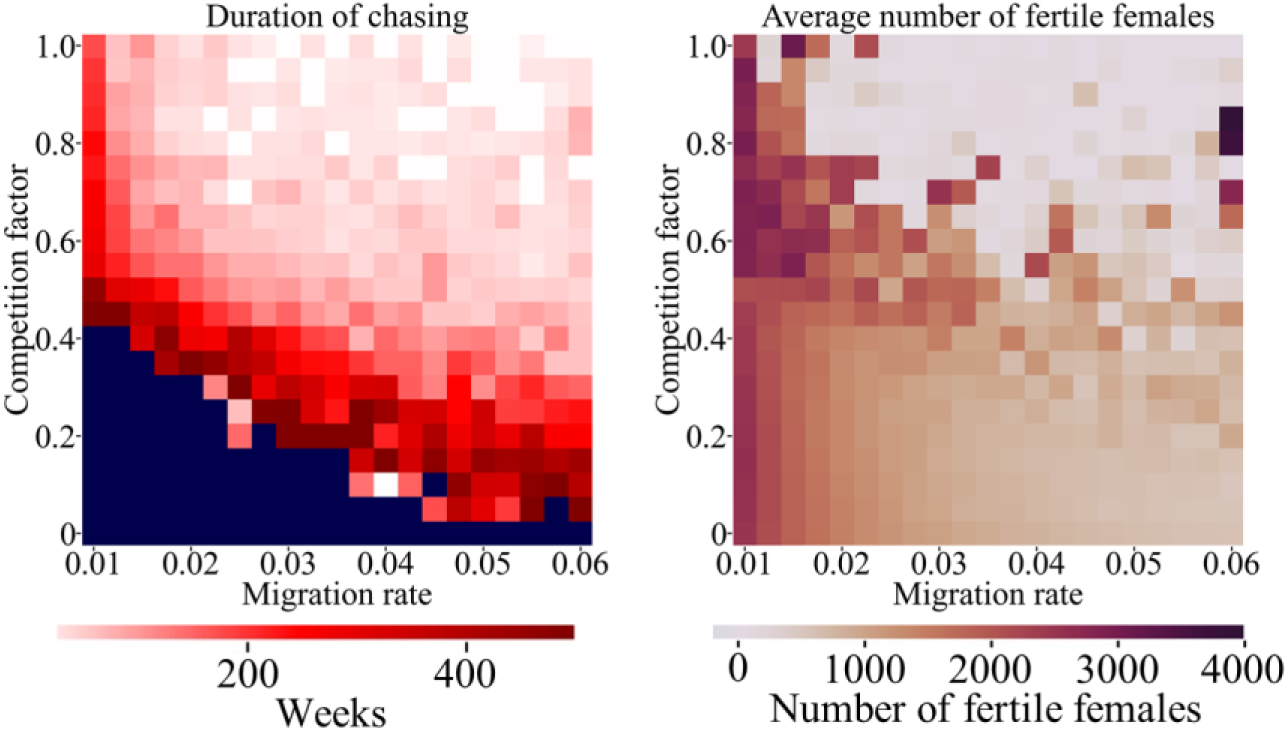
Two-region mosquito model. Drive heterozygotes were released into the middle of a space divided into two regions with two mosquito species with varying migration rate and competition factor. Each species is specialized in one region and competes normally there, while competing at a reduced rate determined by the competition factor in the other region. The duration of chasing and average number of fertile females during chasing is displayed. Blue color refers to points where chasing continued to the end of the simulation in all cases. 20 simulations were assessed for each point in the parameter space.

**Figure S6.**
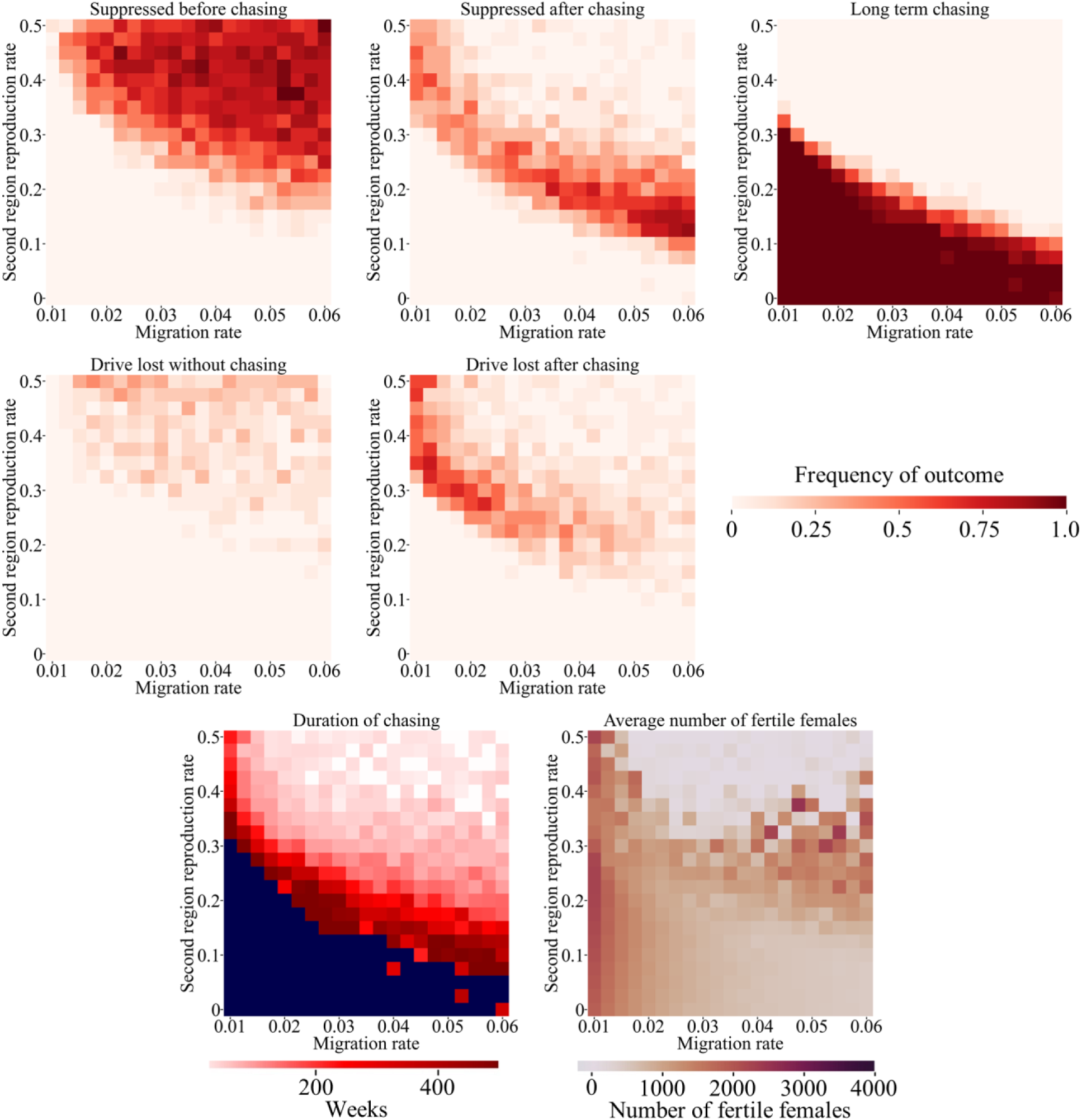
Two-region mosquito model with varying fecundity. Drive heterozygotes were released into the middle of a space divided into two regions with two mosquito species with varying migration rate and competition factor. Each species is specialized in one region and competes normally there, while competing at a reduced rate determined by the competition factor in the other region. The frequency of ultimate outcomes is displayed in the first five panels, and the duration of chasing and average number of fertile females during chasing is displayed in the final two panels. Blue color refers to points where chasing continued to the end of the simulation in all cases. 20 simulations were assessed for each point in the parameter space.

**Figure S7.**
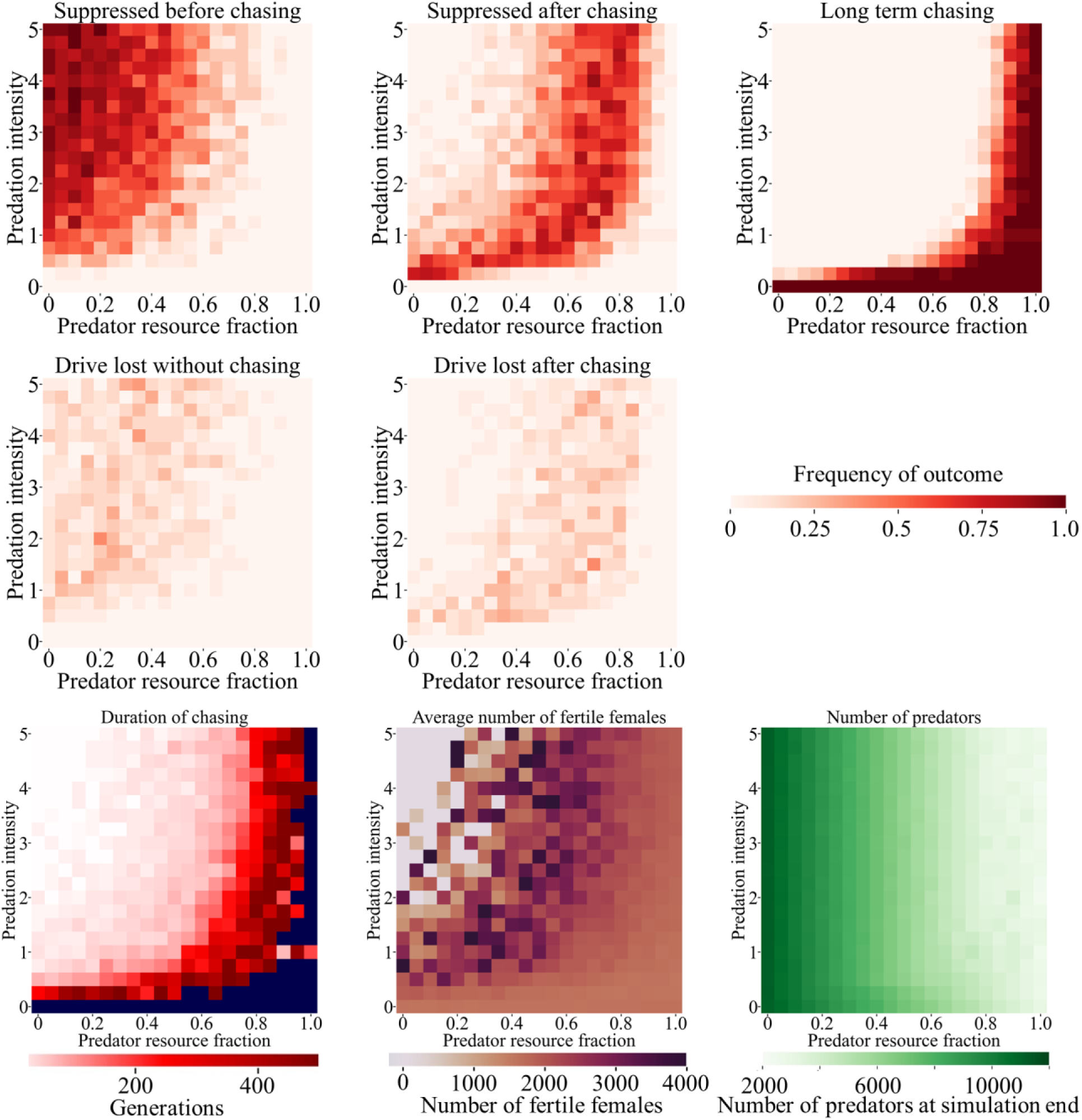
Predator-prey in the discrete-generation model. Drive heterozygotes were released into the middle of a spatial population of 50,000 individuals of the target species that were colocalized with a predator population of 25,000 with varying predation intensity and predator resource fraction (representing the fraction of the importance of the target species to the predator’s diet). The frequency of ultimate outcomes, duration of chasing, average number of fertile females during chasing, and number of predators at the end of the simulation is displayed. Blue color refers to points where chasing continued to the end of the simulation in all cases. 20 simulations were assessed for each point in the parameter space.

**Figure S8.**
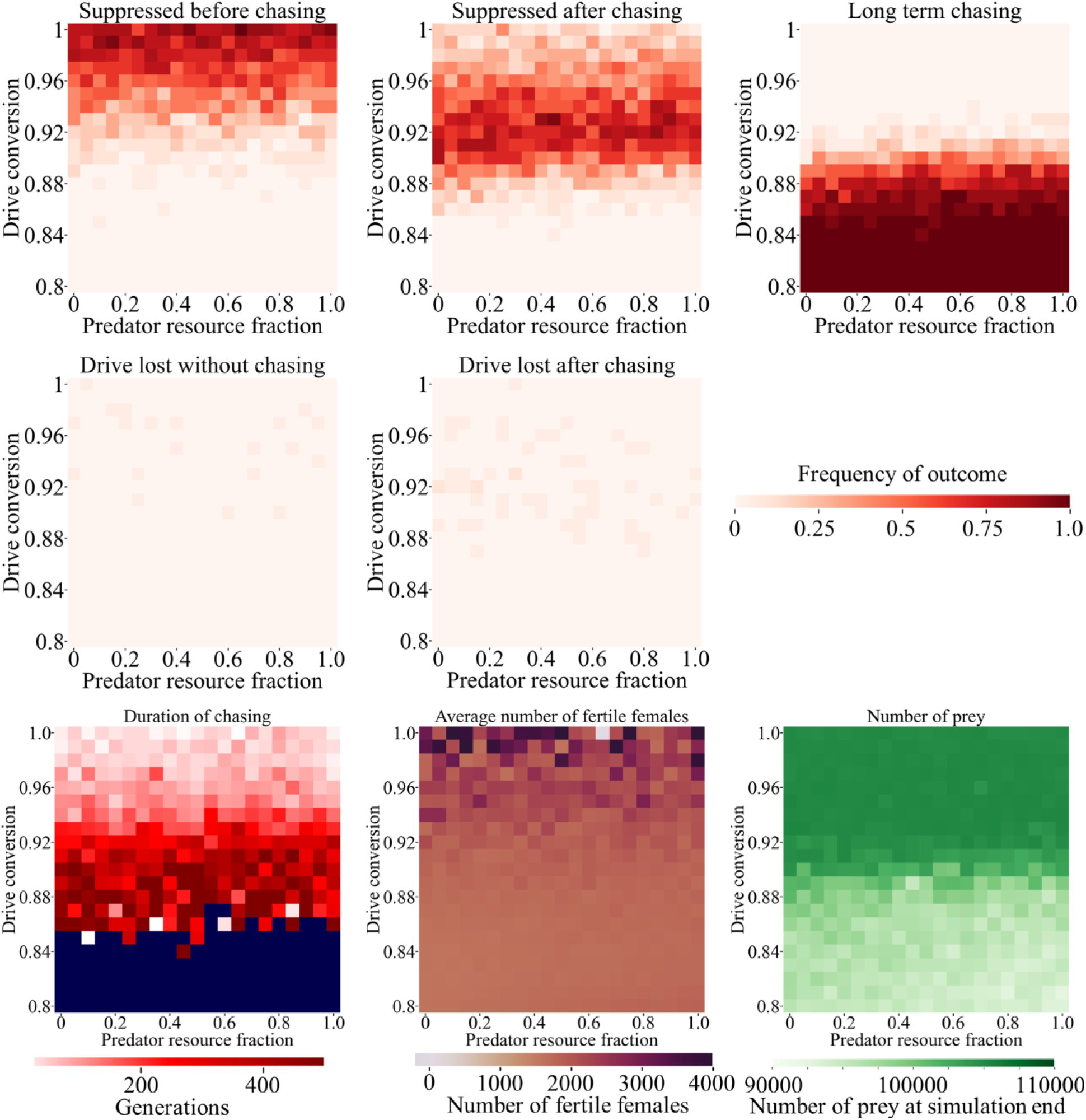
Drive targeting predators. Drive heterozygotes were released into the middle of a spatial population of 50,000 individuals of the target predator species that were colocalized with a prey population of 50,000 with a predation intensity of 1 and varying predator resource fraction (representing the fraction of the importance of the target species to the predator’s diet) and drive conversion rate. The duration of chasing, average number of fertile females during chasing, and number of prey at the end of the simulation is displayed. Blue color refers to points where chasing continued to the end of the simulation in all cases. 20 simulations were assessed for each point in the parameter space.

**Figure S9.**
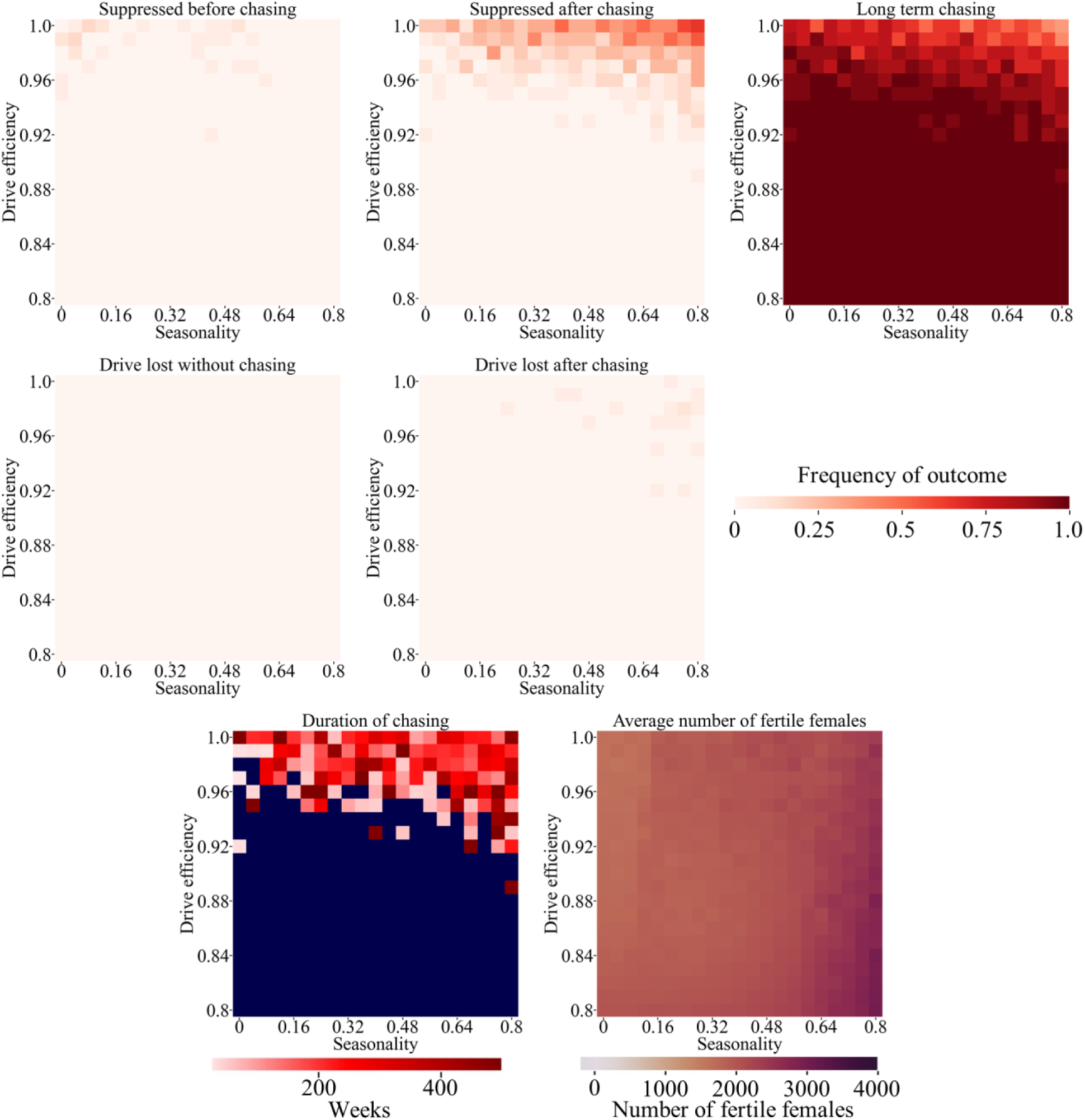
Sharp seasonality. Drive heterozygotes were released into the middle of a mosquito population with 10,000 adult females of the target species and no other species, varying the seasonality factor and drive conversion rate. The population goes through a yearly cycle with the maximum and minimum being above and below the baseline by 10,000 times the seasonality constant and a sharp, linear transition between these lasting for four weeks. Drive outcomes, the duration of chasing, and the average number of fertile females during chasing is displayed. Blue color refers to points where chasing continued to the end of the simulation in all cases. 20 simulations were assessed for each point in the parameter space.

**Figure S10.**
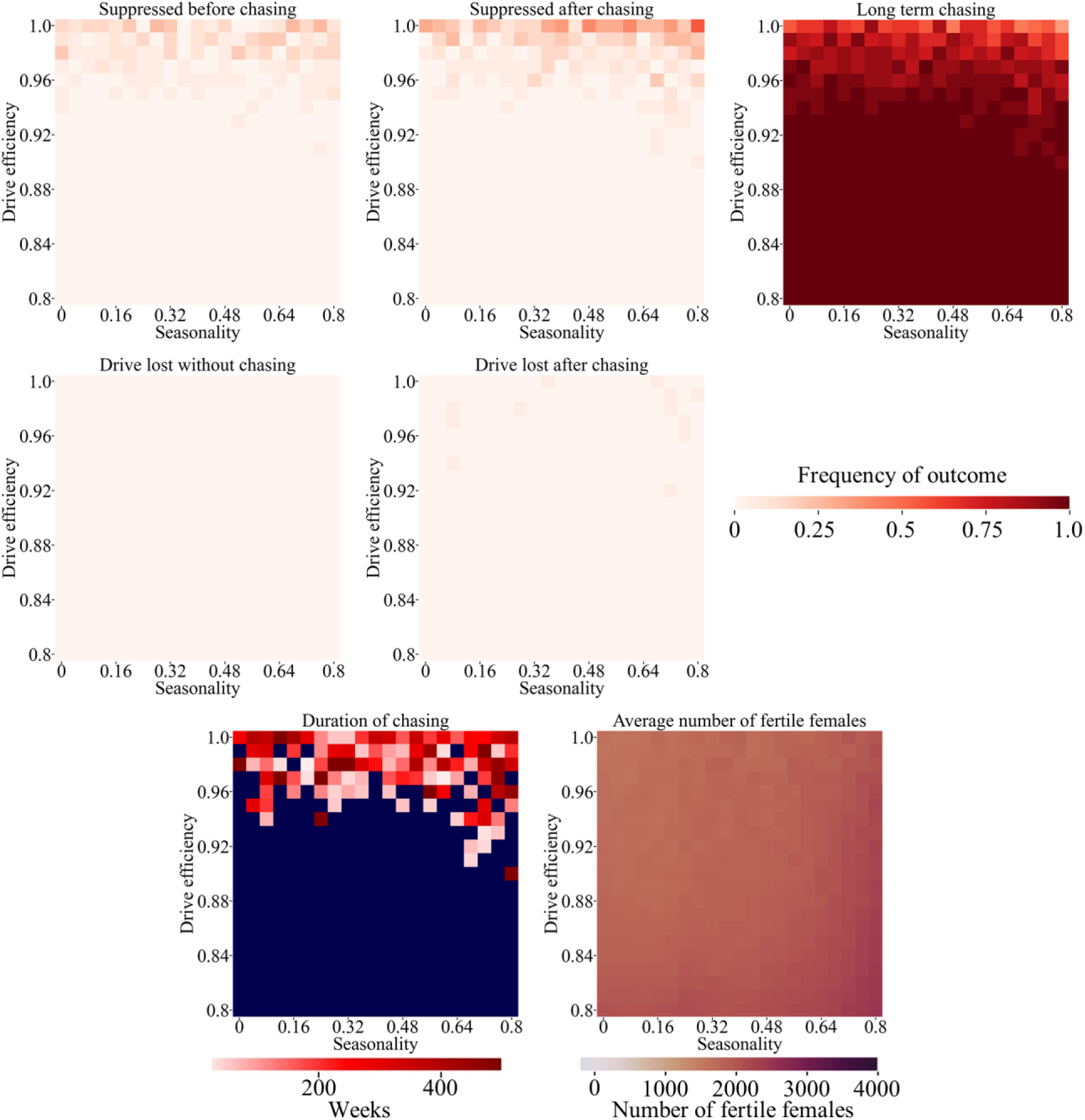
Sinusoidal seasonality. Drive heterozygotes were released into the middle of a mosquito population with 10,000 adult females of the target species and no other species, varying the seasonality factor and drive conversion rate. The population goes through a yearly sinusoidal cycle with the maximum and minimum being above and below the baseline by 10,000 times the seasonality constant. Drive outcomes, the duration of chasing, and the average number of fertile females during chasing is displayed. Blue color refers to points where chasing continued to the end of the simulation in all cases. 20 simulations were assessed for each point in the parameter space.

**Figure S11.**
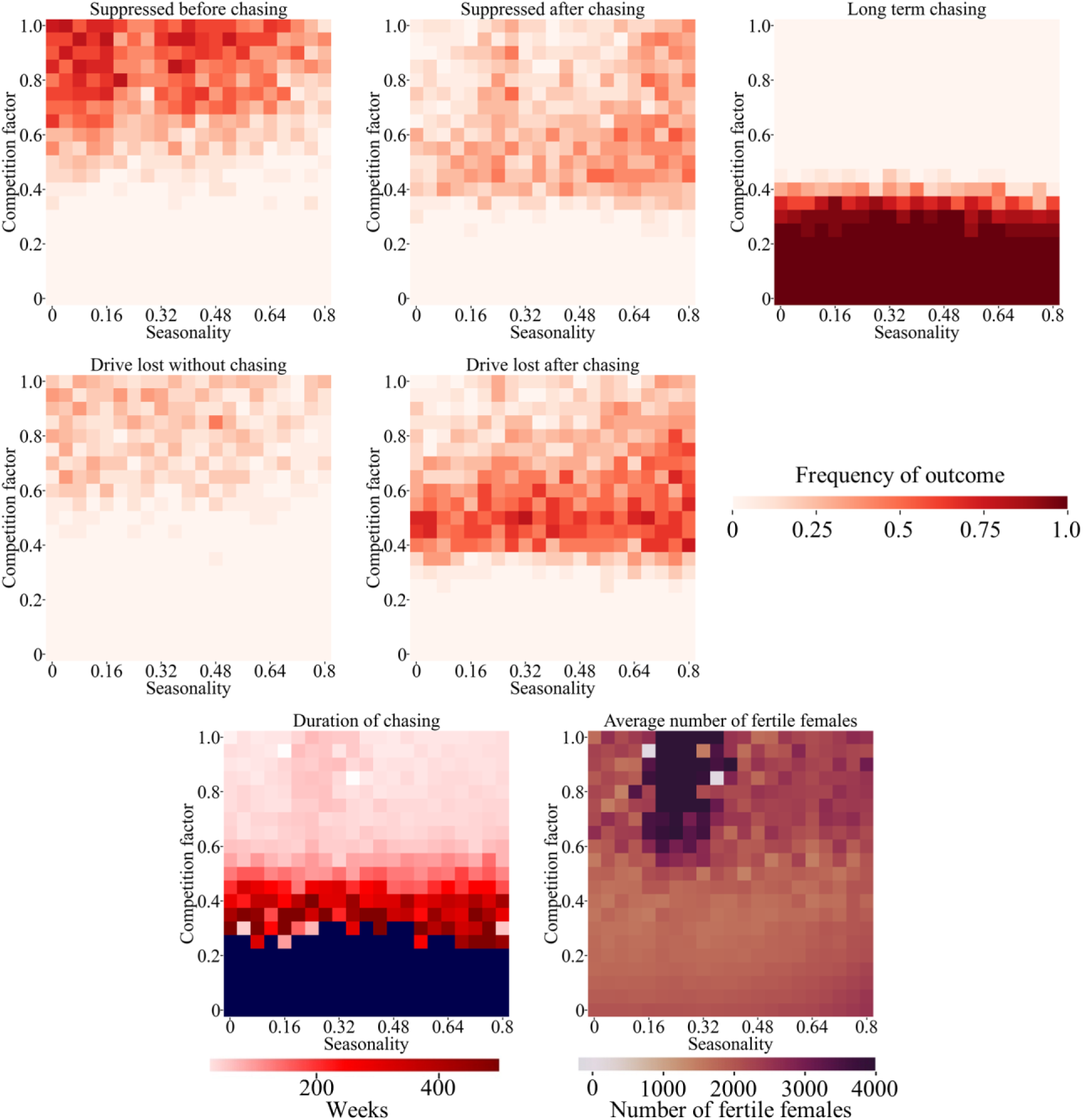
Sinusoidal seasonality and competition. Drive heterozygotes were released into the middle of a mosquito population with a baseline 10,000 adult females of the target species that were colocalized with a competing species with equal population size and a competition factor of 0.5. The population goes through a yearly sinusoidal cycle with the maximum and minimum being above and below the baseline by 10,000 x seasonality constant. The frequency of ultimate drive outcomes, the duration of chasing, and the average number of fertile females during chasing is displayed. Blue color refers to points where chasing continued to the end of the simulation in all cases. 20 simulations were assessed for each point in the parameter space.

**Figure S12.**
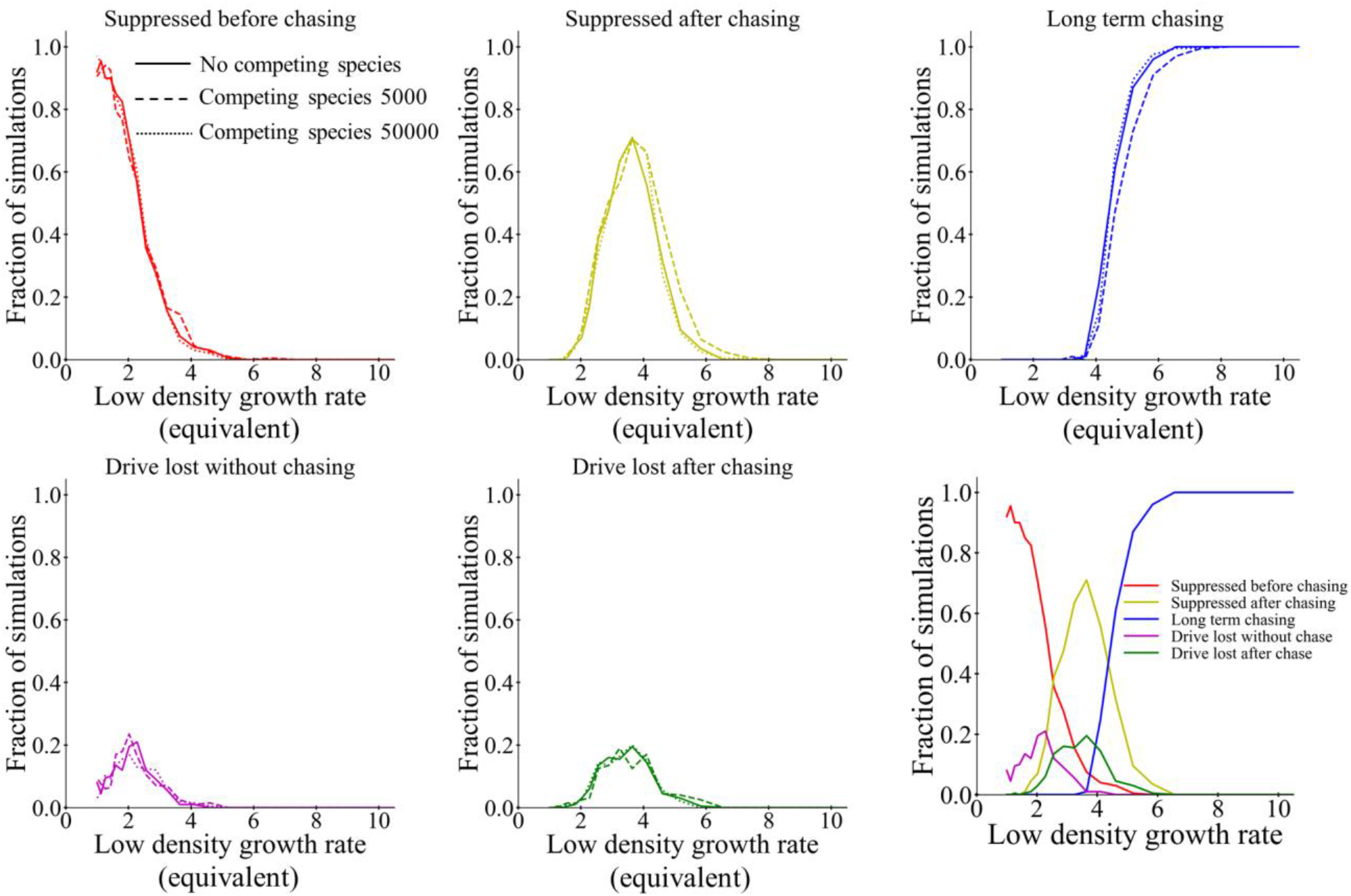
Competition and equilibrium low-density growth rate. Drive heterozygotes were released into the middle of a spatial population of 50,000 individuals of the target species that were alone or were colocalized with a competing species with either 5,000 or 50,000 individuals. The low-density growth rate was varied for the simulations with no competing species and fixed at 10 for the two sets of simulations with a competing species. For simulations with a competing species, the low-density growth rate was fixed at 10, and the competition factor was adjusted match the simulations with no competing species. Specifically, the population of the competing species was assumed to reach its equilibrium value in the absence of the target species (which could occur due to local drive suppression). The growth rate of the target species was then determined under these conditions, assuming that the target species was present at negligible density. This was considered the “equivalent” low-density growth rate. The frequency of ultimate outcomes is displayed for each simulation, and 200 simulations were assessed for each point in the parameter space. The bottom-right panel shows all outcomes together for the simulations with no competing species.

## Notes

https://github.com/jchamper/ChamperLab/tree/main/Suppression-Competition-Modeling

### Competing Interest Statement

The authors have declared no competing interest.

### Summary of Updates

Minor fixes and revisions. Addition of more seasonal modeling data.

